# Stabilizing the CHI3L1-Gal3BP complex remodels the glioblastoma immune microenvironment and synergizes with PD-1 blockade

**DOI:** 10.64898/2026.10.24.761532

**Authors:** Saurabh Upadhyay, Longfei Zhang, Marianna Teplova, Moustafa T. Gabr

**Affiliations:** Department of Radiology, Columbia University Irving Medical Center, New York, NY, 10032, USA; Department of Radiology, Weill Cornell Medicine, New York, NY 10065, USA; Structural Biology Program, Memorial Sloan Kettering Cancer Center, New York, NY 10065, USA

**Author notes:** Corresponding author: Moustafa T. Gabr, PhD.

**Keywords:** glioblastoma, CHI3L1/YKL-40, LGALS3BP, molecular glue, tumor-associated myeloid cells

## Abstract

CHI3L1 (YKL-40) is associated with mesenchymal, treatment-resistant glioblastoma (GBM) and shapes macrophage programs through interactions with galectin-3 or galectin-3-binding protein (Gal3BP). However, the CHI3L1-Gal3BP interaction has not been previously targeted. A surface plasmon resonance screen identified LZ6. Ternary complex formation was assessed by fluorescence-based, biophysical, and mass spectrometry methods. Activity was evaluated in five patient-derived GBM lines, myeloid cultures from eight donors, CRISPR interference with guide-resistant rescue, and a four-arm factorial study of orthotopic GL261 tumors. LZ6 increased Gal3BP engagement 3.4-fold while reducing galectin-3 in vitro. In GBM-conditioned cultures, LZ6 raised phagocytosis 22% to 44% and MHC class II 24% to 39% and lowered CD206 45% to 31% and IL-10 105 to 67 pg/mL, with concordant transcripts. Gal3BP knockdown reduced the response 41%; guide-resistant rescue restored it. Oral LZ6 raised the intratumoral Gal3BP:galectin-3 ratio from 1.02 to 1.56 at unchanged CHI3L1. In tumor tissue LZ6 shifted 20 of 24 immune endpoints, lowering CD206 and IL-10 and raising MHC class II and CD8 effector function. Median survival with LZ6 was 52.4 versus 26.3 days (*P* = 0.0012), and in combination with anti-PD-1 most animals survived to the end of follow-up (median not reached), with a significant factorial interaction (hazard ratio 0.24, 95% CI 0.10-0.57, *P* = 0.0014). LZ6 is a molecular glue of the CHI3L1-Gal3BP complex that engages its target in brain tumor after oral dosing and extends survival alone and synergistically with PD-1 blockade.

## Introduction

Glioblastoma (GBM) remains the most lethal primary malignancy of the adult central nervous system. Maximal resection, radiotherapy and temozolomide produce a median survival close to fifteen months.[1,2] Repeated attempts to extend survival with immune checkpoint blockade have failed in randomized trials,[3] a finding commonly attributed not to a lack of immune cells, but to their immunosuppressive character. GBM is dominated by tumor-associated myeloid cells that suppress, rather than support, antitumor immunity and are actively sustained by soluble factors released by the tumor.[4,5]

Chitinase-3-like protein 1 (CHI3L1), measured clinically as serum or cerebrospinal-fluid YKL-40, is among the most durable correlates of that state.[6] Elevated YKL-40 has been associated with mesenchymal transcriptional identity, invasive behavior, vascular remodeling and poor outcome,[7–9] and CHI3L1 has more recently been identified as a driver within tumor-cell networks that sustain GBM heterogeneity.[10] CHI3L1 is a secreted glycoside hydrolase family 18 protein in which the catalytic glutamate is replaced by leucine; it has no measurable enzymatic activity, and instead acts as an extracellular signaling scaffold through interactions with receptor and co-receptor systems.[11–13]

Critically, the consequences of CHI3L1 signaling depend on which partner it engages. CHI3L1 bound to galectin-3 drives migration and accumulation of protumor, M2-like tumor-associated macrophages, whereas Gal3BP competes with galectin-3 for CHI3L1 and negatively regulates that process; in human GBM, an M1-like macrophage signature is enriched in tumors with low CHI3L1 and high LGALS3BP, and a Gal3BP mimetic peptide reverses immune suppression and slows tumor growth in syngeneic models.[11] In the same work, bulk transcriptomes from sixteen patients treated with nivolumab or pembrolizumab showed higher LGALS3BP in responders than in non-responders, and the authors proposed that interrupting the CHI3L1-galectin-3 complex might synergize with checkpoint inhibition, a prediction that has not been tested with a pharmacological agent.[11] The two partners therefore specify opposite immunological outcomes through the same protein, which makes partner choice, rather than total CHI3L1 activity, the therapeutically meaningful variable.

That architecture is precisely what has kept CHI3L1 outside the reach of conventional pharmacology. There is no catalytic site to inhibit, and the protein’s function is distributed across several partner interactions rather than concentrated in one. Mechanistic work has therefore relied on genetic suppression or neutralizing antibodies,[7,8] and the small molecules reported to date, including high-affinity ligands developed as tool probes, have been characterized principally as inhibitors of CHI3L1-dependent signaling.[14–18] An alternative to inhibition is to change which partner a scaffold protein engages. Small molecules that stabilize rather than block a protein-protein interaction are well established,[19,20] and occupancy of a native ligand pocket is compatible with such a mechanism: rapamycin binds the FKBP12 ligand-binding site and the composite surface recruits mTOR,[21] and auxin acts entirely from within the TIR1 pocket.[22] Applied to CHI3L1, this suggests a strategy in which a groove-binding compound does not silence the protein but redirects it. Because galectin-3 and Gal3BP compete for CHI3L1 and specify opposing macrophage phenotypes,[11] favoring Gal3BP could shift the axis in a direction previously shown to be therapeutically beneficial, while offering pharmacokinetic advantages over peptide-based agents.

Here we identify LZ6, a small molecule identified from a surface plasmon resonance (SPR) screen of a diverse library of small molecules. LZ6 increases CHI3L1 engagement of galectin-3-binding protein (Gal3BP, LGALS3BP) while simultaneously displacing galectin-3, without altering CHI3L1 abundance. We map the composite interface, show that the switch alters myeloid phenotype and function in tumor-conditioned cultures, and demonstrate that an orally administered, tumor-penetrant compound engages the target inside orthotopic GBM, prolongs survival, and synergizes with PD-1 blockade.

## Results

### CHI3L1 and Gal3BP are co-expressed in human GBM

We analyzed TCGA-GBM retrospectively (Broad GDAC Firehose, 28 January 2016). IDH-mutant cases were excluded throughout, since under WHO CNS5 these are astrocytoma, IDH-mutant, grade 4, not GBM. The two Agilent-4502A archives share one patient and were combined by random-effects meta-analysis rather than pooled; the RNAseqV2 cohort shares 135 of its 153 tumors with the arrays and is reported alongside them, not as independent validation. CHI3L1 and LGALS3BP were positively correlated in every cohort, with a meta-analyzed r of 0.42 across the arrays (95% CI 0.34-0.49, *P* = 2 x 10^-22^, n = 488) and r = 0.45 in RNA-seq (*P* = 1 x 10^-8^; Figure 1a). Among the fourteen RNA-seq tumors in no array cohort the correlation was reproduced independently (r = 0.79, *P* = 0.0007). The partners are therefore co-expressed rather than varying independently. Higher CHI3L1 was associated with shorter survival, modestly. Modelled continuously, the meta-analyzed hazard ratio per standard deviation was 1.22 (1.11-1.34, *P* = 4.8 x 10^-5^) with no detectable heterogeneity (Figure 1b), and was weaker and not significant in the smaller, largely overlapping RNA-seq cohort (1.08, 0.89-1.30). Median-based dichotomization, shown for illustration in Figure 1c, did not yield statistically significant results in either array cohort analyzed separately. Accordingly, we describe CHI3L1 as a modest prognostic correlate in IDH-wildtype GBM and do not claim cross-platform validation.

**Figure 1.**
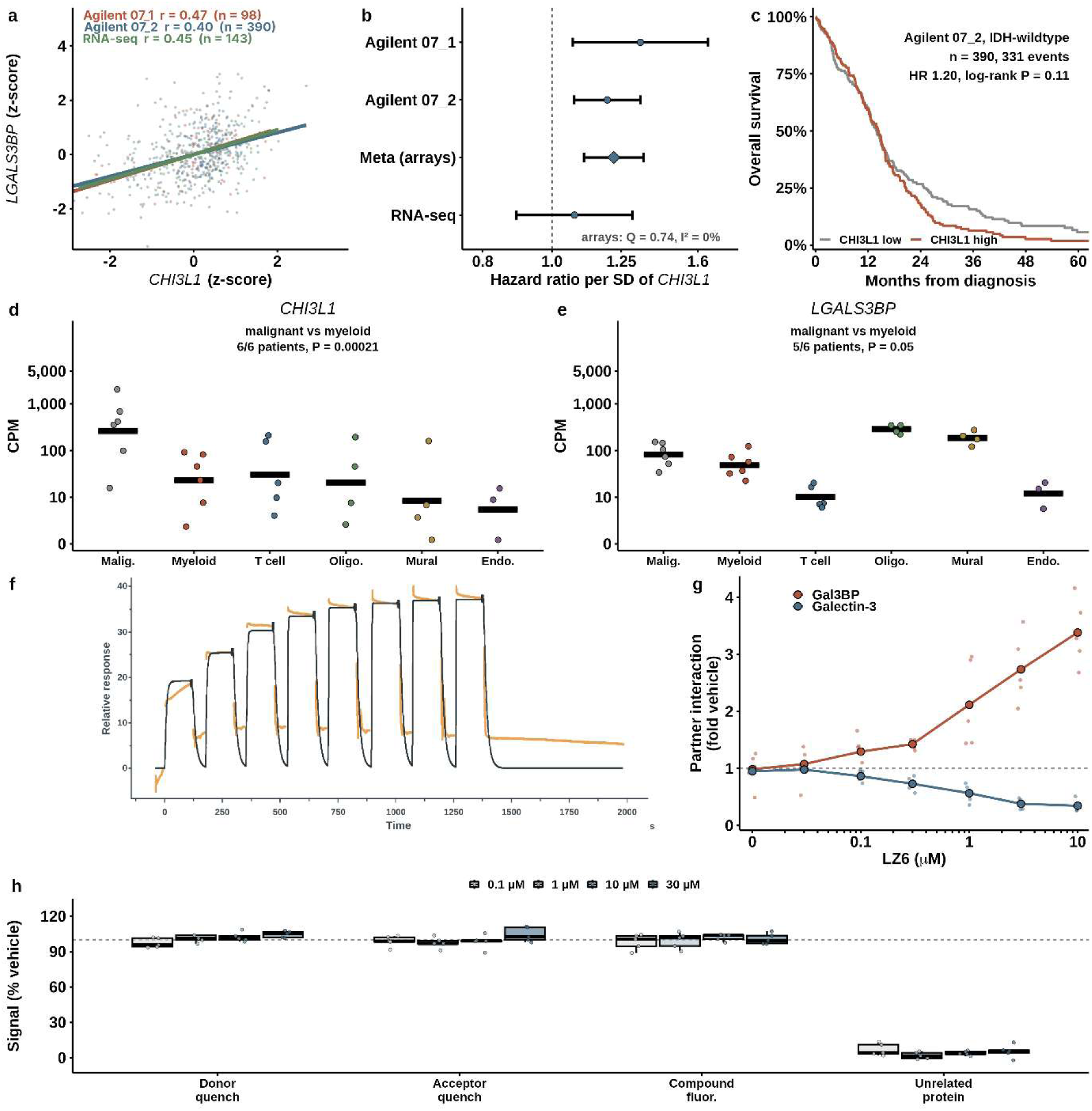
CHI3L1 and Gal3BP in human glioblastoma, and LZ6 as a partner-selective modulator of CHI3L1. (a-c) TCGA-GBM, IDH-wildtype cases only (Broad GDAC Firehose, stddata 28 January 2016). (a) *CHI3L1* against *LGALS3BP*, standardized within cohort, with per-cohort Pearson coefficients. (b) Hazard ratio per standard deviation of *CHI3L1* by cohort, with the random-effects meta-analysis of the two independent array cohorts (diamond); this is the formal survival test. The RNA-seq cohort shares most patients with the arrays and is not independent validation. (c) Kaplan-Meier survival by median *CHI3L1* in the largest array cohort, shown for illustration. (d,e) Compartment expression of *CHI3L1* and *LGALS3BP* in six newly diagnosed glioblastomas (GSE182109); each point is one patient, bars are geometric means, compartments are assigned by canonical lineage markers and cells are aggregated within patient before comparison. (f) Single-cycle kinetics sensorgram for LZ6 binding to immobilized CHI3L1: eight sequential injections of increasing concentration without regeneration between cycles, with the experimental trace in orange and the fitted 1:1 binding model in black. (g) Time-resolved FRET measurement of CHI3L1-Gal3BP and CHI3L1-galectin-3 association across LZ6 concentration; all replicates shown with the mean connected, vehicle at the left-hand tick and the dashed line marking vehicle-equivalent signal. (h) Assay-interference controls to 30 µM; the dashed line marks vehicle-equivalent signal.

To identify the cellular sources of CHI3L1 and Gal3BP, we analyzed single-cell RNA-sequencing data from one region per patient in a published atlas of newly diagnosed GBM.[23] Because cell-type labels were not deposited, we assigned compartments using canonical lineage markers (Supplementary Figure S1). Cells were aggregated by patient and compartment, with the patient serving as the unit of analysis. CHI3L1 expression was predominantly malignant-cell-derived: geometric mean expression was 262 CPM in malignant cells and 22 CPM in myeloid cells, and expression was higher in malignant cells in all six patients (paired t test on log₂ CPM; adjusted *P* = 2.1 × 10^-4^; Figure 1d). LGALS3BP was more broadly expressed, with the highest levels in oligodendrocytes and mural cells (286 and 185 CPM, respectively). Expression in malignant and myeloid compartments was intermediate and similar (81 and 48 CPM, respectively; adjusted *P* = 0.05; Figure 1e). We therefore describe Gal3BP as broadly expressed rather than enriched in a single compartment. The cellular source of a secreted protein need not be the site where it acts. These data identify malignant cells as the predominant source of CHI3L1, consistent with tumor-derived CHI3L1 acting on tumor-associated macrophages.[11]

### LZ6 is a partner-selective modulator of CHI3L1

Screening 4,576 compounds of the Enamine Discovery Diversity Set against immobilized CHI3L1 yielded 53 primary hits at an occupancy threshold of 80% (1.16%), of which 18 (0.39%) survived sensorgram review and advanced to concentration-dependent analysis. LZ6 (Supplementary Figure S2) bound CHI3L1 with a screening SPR binding affinity of 195 nM (Figure 1f). Notably, we confirmed the nanomolar CHI3L1 binding affinity of LZ6 across independent measurements (Supplementary Figure S3). The defining property of LZ6 emerged when the two principal CHI3L1 partner interactions were measured in parallel. Across a 0.03-10 µM range, LZ6 increased CHI3L1-Gal3BP association to 3.4-fold of vehicle while reducing CHI3L1-galectin-3 association to 0.35-fold, in the same assay format and over the same concentrations (Figure 1g). Notably, donor and acceptor quenching and compound autofluorescence controls remained within 89-111% of vehicle at concentrations up to 30 μM, and binding to an unrelated protein did not exceed 13% (Figure 1h; complete control set in Supplementary Figure S4).

### Three orthogonal methods confirm compound-dependent ternary complex formation

Biolayer interferometry (BLI) showed a concentration-dependent increase in CHI3L1-Gal3BP response from 0.77 to 2.52 normalized units (Figure 2a). Native mass spectrometry resolved a ternary species that rose from 6.2% to 66.9% of complex-associated signal (Figure 2b), and SEC-MALS reported higher-mass species rising from 3.6% to 54.4% (Figure 2c). Across both mass-based methods the ternary fraction tracked LZ6 occupancy with a correlation of 0.91 (n = 96; Figure 2e).

**Figure 2.**
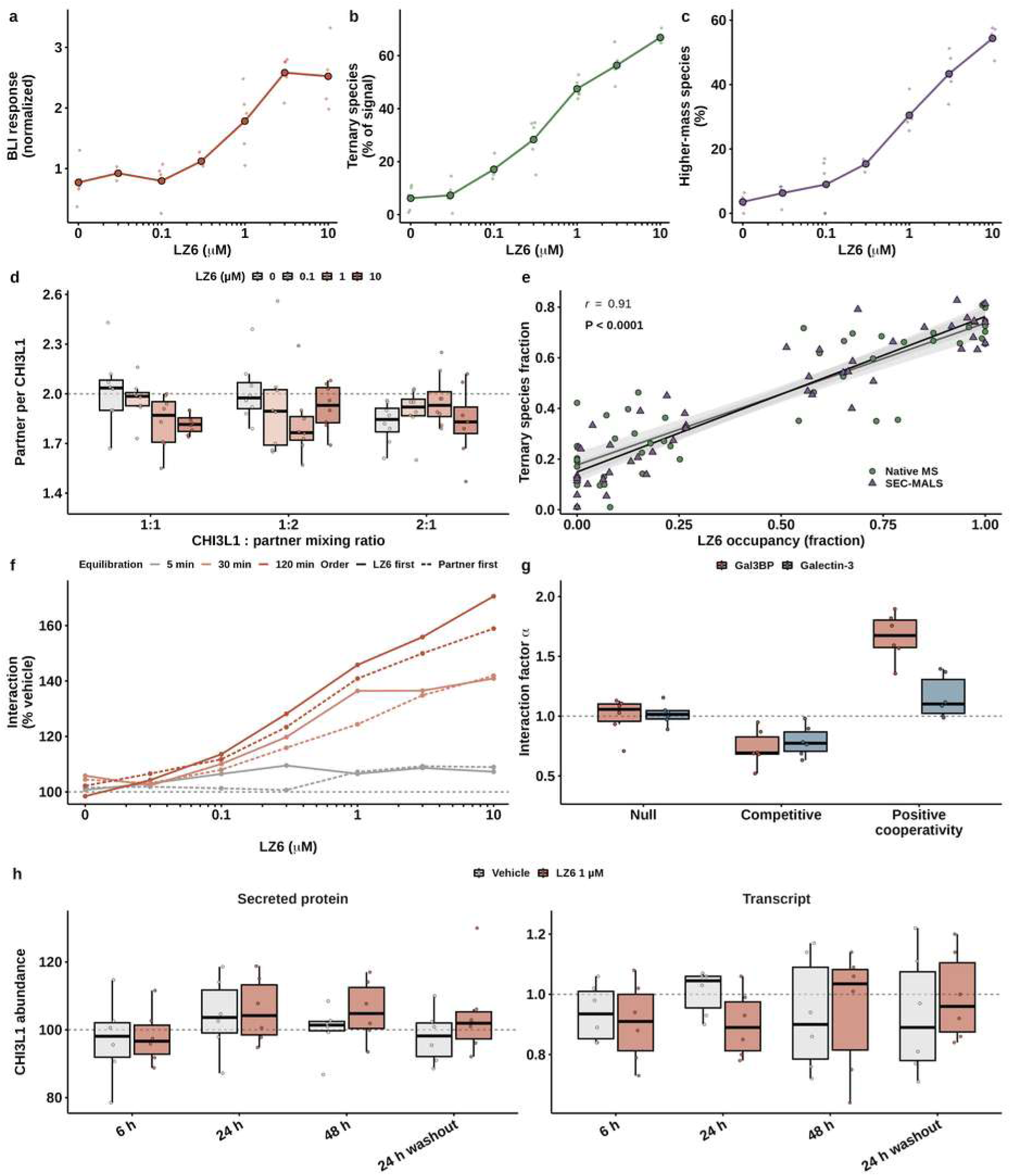
Orthogonal validation of compound-dependent ternary complex formation and its mechanism. (a) Biolayer interferometry, (b) native mass spectrometry ternary species fraction and (c) SEC-MALS higher-mass species across LZ6 concentration; individual replicates shown with means connected. (d) Observed partner-to-CHI3L1 stoichiometry across mixing ratios and compound concentrations; the dashed line marks a ratio of two. (e) Ternary species fraction against LZ6 occupancy for both mass-based methods, with linear fit and Pearson correlation. (f) Effect of order of addition and equilibration time. (g) Fitted interaction factor for null, competitive and positively cooperative models, by partner, across six independent protein preparations. (h) CHI3L1 secreted protein and transcript during and after LZ6 exposure.

The observed stoichiometry was approximately two partner molecules per CHI3L1 and did not change with CHI3L1:partner mixing ratio or with compound concentration (Figure 2d), indicating that LZ6 increases the proportion of complex rather than altering its composition. The effect was independent of the order in which compound and partner were added but increased with equilibration time, from 107% of vehicle at 5 minutes to 171% at 120 minutes (Figure 2f), a pattern consistent with a shift in equilibrium rather than kinetic trapping. Fitting the titrations to null, competitive and positively cooperative models returned an interaction factor of 1.66 +/- 0.20 for Gal3BP against 1.16 +/- 0.18 for galectin-3 (Figure 2g); model selection criteria for every preparation are given in Supplementary Figure S5, and the full stoichiometry series in Supplementary Figure S6. Neither CHI3L1 transcript nor secreted CHI3L1 protein changed at 6, 24 or 48 hours of exposure, or after washout (Figure 2h). The compound therefore acts on the interaction rather than on the abundance of the target.

### LZ6 promotes a composite CHI3L1-Gal3BP interface

To define the molecular basis of LZ6 recognition, we determined the crystal structure of the human CHI3L1-LZ6 complex at 2.75 Å resolution (PDB ID: 10JZ). The structure places LZ6 lengthwise within the glycoside hydrolase family 18 groove, enclosed by an aromatic environment including Trp99, Tyr206 and Trp352, with one end of the ligand oriented toward the solvent-accessible entrance of the pocket near Glu290 (Figure 3a-d). The bound ligand occupies a predominantly neutral-to-negatively charged region of the groove, with its distal end directed toward solvent (Supplementary Figure S7). Data collection and refinement statistics are given in Supplementary Table S1.

**Figure 3.**
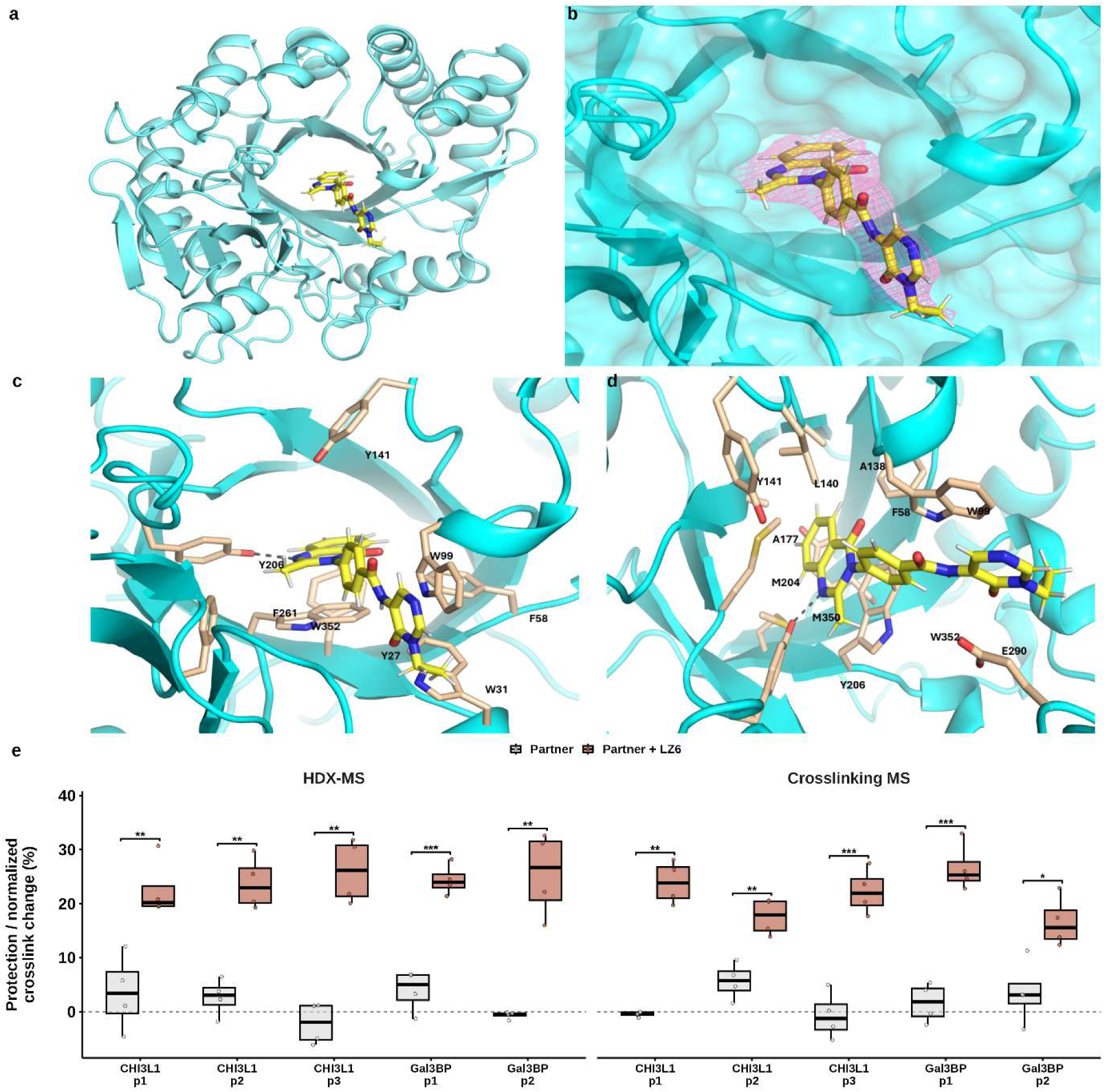
A composite CHI3L1-Gal3BP interface. (a) Overall fold of the CHI3L1-LZ6 complex (PDB 10JZ) with LZ6 in yellow. (b) Electron density for the bound ligand within the carbohydrate-binding groove. (c,d) Aromatic and aliphatic residues lining the groove, in two orientations. (e) Protection by hydrogen-deuterium exchange and crosslinking mass spectrometry across monitored CHI3L1 and Gal3BP peptides, with and without LZ6; four independent preparations per condition, compared by Welch t test with Benjamini-Hochberg adjustment across the ten regions. The exact Wilcoxon rank-sum floor at four versus four is P = 0.0286, which every separated region would return identically, so the rank test was not used.

Hydrogen-deuterium exchange and crosslinking mass spectrometry were used to test whether LZ6 in the groove creates a surface that the partner also contacts. With partner alone, protection across the monitored peptides was near baseline (1.5 ± 4.6% by HDX-MS and 2.0 ± 4.3% by crosslinking). Adding LZ6 raised protection to 24.5 ± 5.1% and 21.4 ± 5.4%, and protection increased in every monitored region of both CHI3L1 and Gal3BP by both methods (Figure 3e).

### Partner switching alters tumor-cell behavior and depends on the CHI3L1-LGALS3BP axis

LZ6 reduced 3D spheroid viability across five patient-derived GBM lines and a CHI3L1-expressing GL261 line, reaching approximately 72% of vehicle at 30 µM, with comparable effects in tumor-only and myeloid co-culture spheroids (Figure 4a,b). The magnitude of direct cytotoxicity was modest across a thousand-fold concentration range. Lactate dehydrogenase release and post-washout sphere-forming capacity followed the same concentration dependence (Figure 4c,d). Normal human astrocytes were comparatively spared, retaining 90.0% viability at 30 µM against 72.2% for GBM lines (n = 6 astrocyte culture replicates versus 6 line means, *P* = 2.5 x 10^-5^; Figure 4e). Target engagement was detectable in the cellular setting: the CHI3L1-Gal3BP complex index in conditioned media rose in a concentration-dependent manner across all six models (Figure 4f). Baseline target expression across models is given in Supplementary Figure S8.

**Figure 4.**
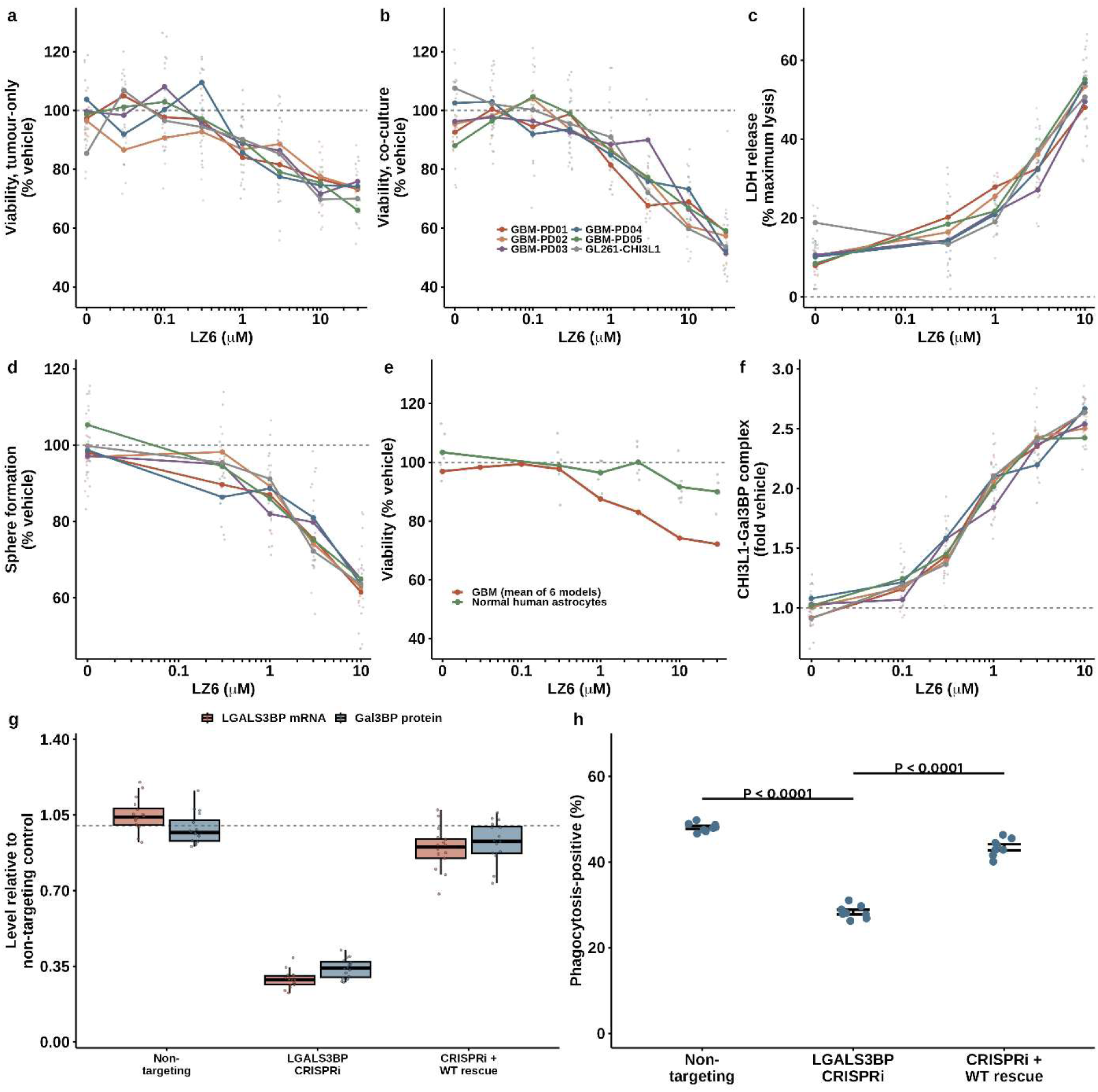
Partner switching reduces tumor-cell viability and depends on the CHI3L1-LGALS3BP axis. (a,b) Three-dimensional spheroid viability across five patient-derived glioblastoma lines and a CHI3L1-expressing GL261 line, in tumor-only and myeloid co-culture settings. (c) Lactate dehydrogenase release. (d) Post-washout sphere-forming capacity. (e) Normal human astrocyte viability compared with the mean of six glioblastoma models. (f) CHI3L1-Gal3BP complex index in conditioned media. (g) LGALS3BP transcript and secreted Gal3BP protein across CRISPR interference genotypes; a guide-resistant wild-type construct restores both to approximately 90% of the non-targeting control. (h) Phagocytosis across the same genotypes in eight donors; each point is one donor and comparisons are donor-paired t tests. Silencing LGALS3BP reduces the response and the guide-resistant construct restores it.

CRISPR interference reduced LGALS3BP transcript to 29% and secreted Gal3BP to 34% of baseline, and a CRISPRi-resistant wild-type construct restored them to 90% and 93% (Figure 4g; per-donor values in Supplementary Figure S9). The functional results paralleled the expression data: the proportion of phagocytic cells decreased from 48.1% to 28.4% following knockdown (*P* = 8.8 × 10^-9^) and was partially restored to 43.5% by the guide-resistant construct (P = 2.5 × 10^-7^ vs. knockdown; Figure 4h). Because the resistant construct restores expression and largely restores function, the phagocytic response depends on Gal3BP itself rather than on an off-target effect of the guides.

### Tumor conditioning amplifies a coordinated myeloid phenotype shift

In GBM-conditioned myeloid cultures from eight independent donors, LZ6 produced concentration-dependent changes in every endpoint measured. Phagocytosis rose from 21.6% to 44.2% of cells and MHC class II-positive cells from 24.0% to 39.1%, while CD206-positive cells fell from 45.2% to 31.0% and secreted IL-10 from 105 to 67 pg/mL, each across 0 to 5.71 µM and each monotonic with concentration (all adjusted *P* < 10^-5^; Figure 5a-d).

**Figure 5.**
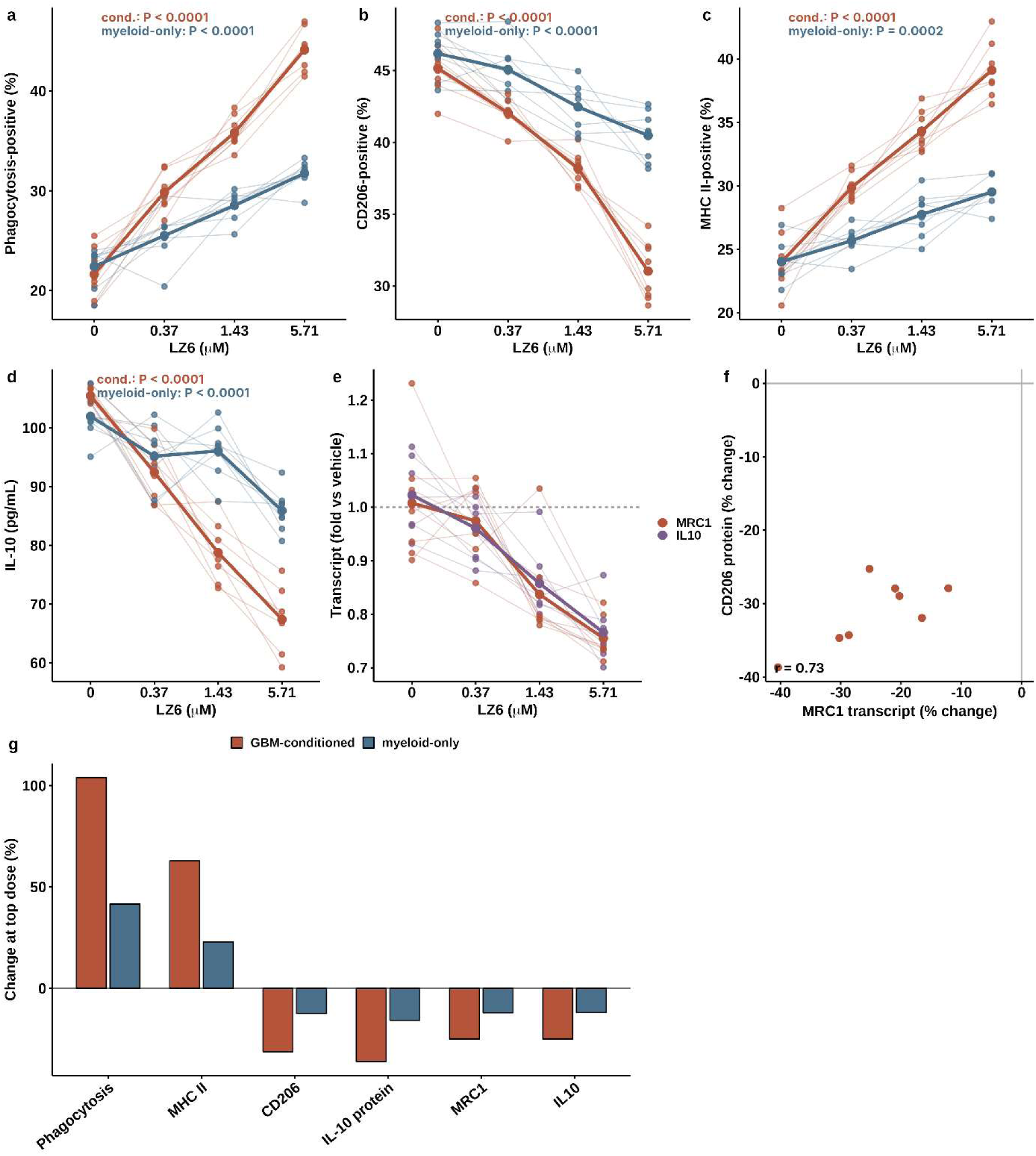
A coordinated myeloid phenotype shift, amplified by tumor conditioning. Throughout, the biological unit is the independent donor (n = 8); technical replicates were averaged within donor before analysis, plotted points are donor means, and comparisons are donor-paired t tests with Benjamini-Hochberg adjustment within endpoint. (a) Phagocytosis, (b) CD206-positive fraction, (c) MHC class II-positive fraction and (d) secreted IL-10 across LZ6 concentration in GBM-conditioned and myeloid-only cultures, with donors connected and bold lines showing group means; adjusted P values at the top concentration are shown for each context. (e) MRC1 and IL10 transcripts in GBM-conditioned cultures, relative to matched vehicle. (f) Per-donor change in MRC1 transcript against change in CD206 surface protein at the top concentration. (g) Change at the top concentration for all six endpoints in both assay contexts.

Tumor conditioning set the magnitude of these changes but not their direction. In myeloid-only cultures every endpoint moved the same way as in conditioned cultures and remained significant, but the effect was roughly a third to a half as large: phagocytosis rose 42% against 104%, MHC class II 23% against 63%, and CD206 fell 12% against 31% (Figure 5g). The tumor-conditioned environment therefore amplifies a response that the compound produces in myeloid cells on its own, rather than being required for it.

Transcript and protein measurements agreed. MRC1 fell to 0.76 and IL10 to 0.77 of vehicle in the same donors and at the same concentrations at which CD206 surface protein and secreted IL-10 fell, all samples collected at a single matched timepoint (Figure 5e). Across donors, the change in MRC1 transcript tracked the change in CD206 protein (r = 0.73; Figure 5f). The coordinated movement of a surface marker, its transcript, a secreted cytokine, its transcript, an antigen-presentation marker and a functional phagocytic readout in the same direction is what distinguishes a phenotypic shift from an isolated endpoint effect. The direction of the surface-marker changes in tumor-conditioned cultures, with CD206 falling and CD86 and MHC class II rising, matches the phenotype previously obtained by competing galectin-3 off CHI3L1 with a Gal3BP mimetic peptide.[11] Tumor-cell migration and invasion decreased with concentration across all models, reaching approximately 65-70% of vehicle at 10 µM (Supplementary Figure S10).

### Oral LZ6 reaches intracranial tumor, engages CHI3L1-Gal3BP and prolongs survival

Oral administration of LZ6 produced dose-dependent plasma exposure across 3, 10 and 30 mg/kg, with an AUC-versus-dose slope of 0.80 (95% confidence interval 0.74-0.86), indicating slightly less than dose-proportional exposure (Figure 6a,b; full profiles in Supplementary Figure S11). The compound showed acceptable solubility and intrinsic clearance, a hERG inhibition margin, and efflux ratios near three that were abolished by selective transporter inhibition (Supplementary Figures S12 and S13). In animals bearing intracranial tumors, paired plasma, normal brain and tumor sampling showed tumor concentrations exceeding those in normal brain at every timepoint (Figure 6c). Unbound partition coefficients rose over time and were higher in tumor than in normal brain from four hours onward, reaching 0.61 in tumor against 0.22 in normal brain at eight hours (*P* = 0.002 and 0.004 at 4 and 8 hours; Figure 6d). Fourteen days of daily dosing produced no weight loss relative to vehicle (Figure 6e); individual trajectories, clinical scores and the remaining satellite pharmacodynamic endpoints are given in Supplementary Figure S14.

**Figure 6.**
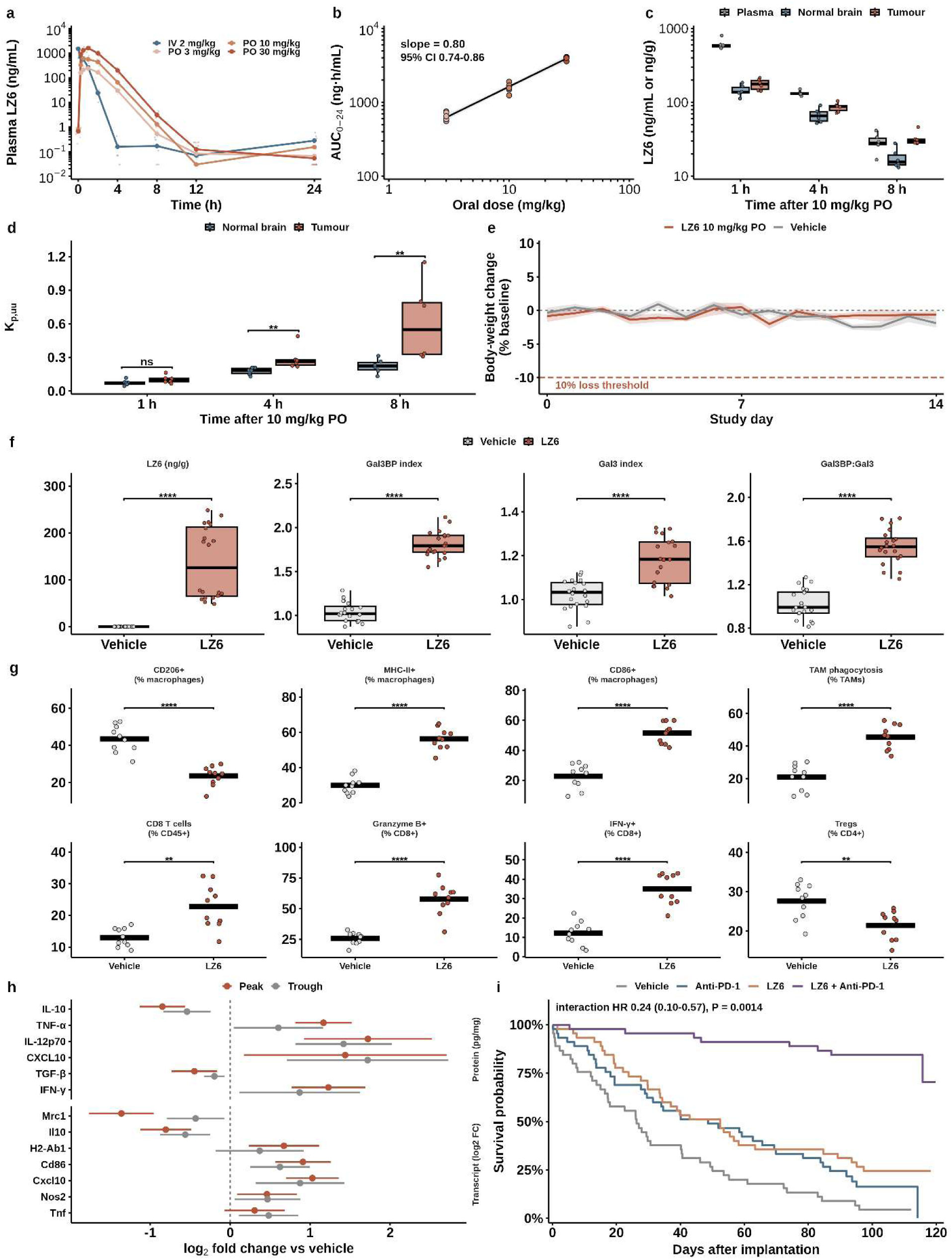
Oral exposure, tumor penetration, in vivo target engagement and survival. (a) Plasma concentration-time profiles following intravenous and oral administration. (b) Dose proportionality of oral exposure with fitted slope and confidence interval. (c) Paired plasma, normal brain and intracranial tumor concentrations after 10 mg/kg orally. (d) Unbound partition coefficients for normal brain and tumor. (e) Body-weight change during fourteen days of daily dosing; the dashed line marks a 10% loss threshold. (f) Tumor drug concentration, CHI3L1-Gal3BP complex index, CHI3L1-galectin-3 complex index and their per-animal ratio. The two complex indices are assay-specific findings in tumor homogenate; the ratio was computed for each animal from its own paired measurements and tested at the animal level. (g) Intratumoral myeloid and lymphoid endpoints by flow cytometry at peak sampling; each point is one animal (n = 10 per arm), bars are means, and comparisons are Welch t tests with Benjamini-Hochberg adjustment across all twenty-four endpoints within phase. (h) Soluble mediators and transcripts as log2 fold change over vehicle at both sampling phases, with 95% confidence intervals. Three below-limit-of-quantification records, one vehicle animal each in IL-12p70, CXCL10 and interferon-gamma, were set to half the lowest observed non-zero value for that analyte. (i) Kaplan-Meier survival for orthotopic GL261 tumours, four arms of 45 animals (180 animals, 123 events); the factorial interaction term is shown on the panel. Log-rank P values against vehicle are 0.014 for anti-PD-1, 0.0012 for LZ6 and below 10-10 for the combination.

In tumor-bearing animals, LZ6 reached 64 ng/g in tumor at trough and 209 ng/g at peak, with no compound detectable in vehicle animals at either sampling point. The intratumoral CHI3L1-Gal3BP complex index rose from 1.02 to 1.72 at trough and to 1.90 at peak (both *P* < 10^-10^), and tumor drug concentration correlated with the Gal3BP index across treated animals (r = 0.63, *P* = 0.003), linking exposure to engagement. The CHI3L1-galectin-3 index also rose in tumor, by much less, from 1.02 to 1.14 at trough and 1.23 at peak; this is an assay-specific finding in tumor homogenate and is reported separately from the reduction in galectin-3 association seen in the purified assay. The ratio of the two indices was computed for each animal from its own paired measurements and tested at the animal level, rising from 1.02 to 1.53 at trough and 1.56 at peak (*P* = 4.0 x 10^-6^ and 1.6 x 10^-8^, n = 10 per arm per phase; Figure 6f). Total CHI3L1 abundance was unchanged (*P* = 0.63), so engagement occurs without alteration of target levels. Engagement was accompanied by a broad shift in the tumor immune compartment, profiled across twenty-four endpoints in ten animals per arm at each sampling phase.

Total macrophage abundance was unchanged (40.6% against 38.4% of CD45-positive cells, adjusted *P* = 0.34), but their phenotype was not. CD206-positive macrophages fell from 43.5% to 23.6% while MHC class II-positive macrophages rose from 29.8% to 56.3% and CD86-positive macrophages from 22.8% to 51.5%, and ex vivo phagocytic capacity of tumor-associated macrophages rose from 21.1% to 45.5% (all adjusted *P* < 10^-4^ at peak; Figure 6g). The lymphoid compartment moved in the same direction: CD8 T cells rose from 13.0% to 22.8% of CD45-positive cells (adjusted *P* = 0.0028), and among them granzyme B-positive cells rose from 25.9% to 57.7% and interferon-gamma-positive cells from 12.2% to 35.0% (both adjusted *P* < 10^-4^), while regulatory T cells fell from 27.6% to 21.4% of CD4^+^ T cells (adjusted *P* = 0.0059). Neutrophils and Ly6C-positive monocytes did not change. Soluble mediators and transcripts agreed with the cellular readouts at both sampling phases (Figure 6h): IL-10 fell 0.85 log2 units and TGF-beta 0.45, while TNF-alpha rose 1.17, IL-12p70 1.72, CXCL10 1.44 and interferon-gamma 1.23 at peak; Mrc1 and Il10 transcripts fell 1.36 and 0.81 log2 units while H2-Ab1, Cd86 and Cxcl10 rose 0.67, 0.91 and 1.03. The directions observed in donor myeloid cultures therefore reproduce in tumor tissue, with the addition of CD8 effector function the cultures could not report.

In orthotopic GL261 tumors randomized across four arms of 45 animals each (180 animals), median survival was 26.3 days with vehicle, 52.4 days with LZ6 (log-rank *P* = 0.0012) and 48.7 days with anti-PD-1 (*P* = 0.014). Median survival was not reached in the combination arm, in which only 8 of 45 animals had died at the end of follow-up (*P* < 10^-10^ against vehicle; Figure 6i), and the combination exceeded each monotherapy (*P* = 1.0 x 10^-8^ against LZ6 alone and 5.9 x 10^-12^ against anti-PD-1 alone). In the factorial Cox model each agent contributed a significant main effect (LZ6 hazard ratio 0.45, 95% CI 0.29-0.72, *P* = 7.4 x 10^-4^; anti-PD-1 hazard ratio 0.58, 95% CI 0.38-0.91, *P* = 0.017), and the interaction term was itself significant (hazard ratio 0.24, 95% CI 0.10-0.57; Wald *P* = 0.0014). Proportional hazards were satisfied for every term and globally (global *P* = 0.94). The combination therefore acts synergistically rather than additively: multiplicative additivity predicts a combination hazard ratio of 0.27, and the observed value is 0.06.

## Discussion

LZ6 changes which partner CHI3L1 preferentially engages, increasing association with Gal3BP while displacing galectin-3, with no change in transcript or protein level in culture or in tumor tissue. The direction of that switch is not arbitrary. Galectin-3 and Gal3BP compete for CHI3L1, and the two complexes specify opposing macrophage programs: the galectin-3 complex promotes protumor M2-like macrophage migration and accumulation, whereas Gal3BP engagement opposes it. A Gal3BP mimetic peptide reverses immune suppression in syngeneic GBM.[11] LZ6 moves the axis in that same direction.

Because galectin-3 and Gal3BP compete for CHI3L1, stabilizing one complex may disfavor formation of the other. Thus, the therapeutic effect could reflect both increased CHI3L1-Gal3BP association and decreased CHI3L1-galectin-3 association; the present data do not distinguish these contributions. The opposing interaction changes are a key observation: in the same assay format, LZ6 increased CHI3L1-Gal3BP association 3.4-fold while reducing CHI3L1-galectin-3 association to 0.35-fold. These opposing effects argue against a nonspecific mechanism such as aggregation, assay interference, or avidity. Three orthogonal methods, two of them mass-based and label-free, reproduced the ternary species. The invariant stoichiometry across mixing ratios indicates that the compound changes the proportion of complex rather than its composition, and the dependence on equilibration time but not on order of addition is what would be expected of a thermodynamic shift.

The X-ray co-crystal structure of CHI3L1 with LZ6 places the compound in the carbohydrate-binding groove. Compound-dependent protection of peptides from both CHI3L1 and Gal3BP, which was near baseline when the proteins were incubated without compound, supports a composite interface. Because the peptides were not mapped to specific residues, the interface remains unresolved; determining whether Gal3BP contacts LZ6 directly or engages a remodeled surface will require a ternary structure or residue-level mapping. Together, these findings support partner-selective, positively cooperative stabilization of the CHI3L1-Gal3BP complex. Three orthogonal methods detected compound-dependent ternary-complex formation, and the ternary fraction scaled with CHI3L1 occupancy.

This study adds an orally administered small molecule, an X-ray co-crystal structure, evidence for a composite interface, and direct measurement of tumor target engagement without changes in CHI3L1 abundance. The immune changes corroborate the therapeutic direction established by Chen and colleagues; they do not establish a new mechanism. In the presence of LZ6, GBM spheroid viability decreased to approximately 72% of vehicle, with relative sparing of normal astrocytes. Its larger effects were in the myeloid compartment and were consistent across eight donors. Phagocytosis and MHC class II increased, while CD206, secreted IL-10, and MRC1 and IL10 transcripts decreased. Tumor conditioning amplified these responses but was not required; effects in myeloid-only cultures were approximately one-third to one-half those in conditioned cultures. The phagocytic response depended on Gal3BP: LGALS3BP silencing reduced it by 41%, while a guide-resistant wild-type construct that restored expression to 90% of control largely rescued function.

The in vivo results are consistent with the proposed mechanism. An orally administered LZ6 reaches intracranial tumor at concentrations exceeding those in normal brain, with unbound partitioning rising over time and favoring tumor from four hours onward. In treated tumors the CHI3L1-Gal3BP complex index rises from 1.02 to 1.90 at peak while total CHI3L1 is unchanged, and the index tracks tumor drug concentration across animals, which is target engagement of the intended kind. The survival result is strong in both its main effects and its interaction. In a four-arm study of 180 animals, LZ6 alone extended median survival from 26.3 to 52.4 days (*P* = 0.0012) and anti-PD-1 alone to 48.7 days (*P* = 0.014); median survival was not reached in the combination arm, where only 8 of 45 animals had died at the end of follow-up, and the combination exceeded each monotherapy by a wide margin. In the factorial model, each agent had a significant main effect, and their interaction was significant (hazard ratio, 0.24; 95% CI, 0.10-0.57; Wald *P* = 0.0014). Proportional-hazards assumptions were met for each term and globally (*P* = 0.94). The combination therefore exceeded the effect predicted by multiplicative additivity: the predicted hazard ratio was 0.27, compared with an observed value of 0.06.

Immune profiling suggests a possible mechanism. LZ6 increased CD8^+^ T-cell infiltration and the proportion of CD8^+^ T cells expressing granzyme B and interferon-γ, while reducing regulatory T cells and TGF-β and increasing IL-12p70 and CXCL10. These changes could improve antitumor immunity, but the immune and survival endpoints were measured in separate cohorts, and the combination arm was not profiled. Thus, the immune changes have not been established as mediators of the survival benefit. Efficacy was tested in GL261, which has limitations as a model of human GBM; patient-derived studies were limited to cell culture.

## Materials and Methods

### Human cohort analyses

TCGA-GBM expression and clinical data were obtained from the Broad Institute GDAC Firehose (stddata_2016_01_28) as two Agilent-4502A array cohorts and one RNAseqV2 cohort, restricted to primary tumors with IDH-mutant cases excluded. CHI3L1 was standardized within cohort and modelled continuously by Cox proportional hazards, and cohort estimates were combined by random-effects meta-analysis. Single-cell data for six newly diagnosed GBM, one region per patient, were obtained from GSE182109; because compartment labels were not deposited, cells were assigned by canonical lineage marker scores and aggregated within patient, with the patient as the unit of analysis. Filtering criteria, marker sets and software versions are given in Supplementary Methods.

### Protein production, SPR screening and validation

Human CHI3L1 (residues 1-383, C-terminal HRV 3C site and polyhistidine tag) was expressed in Expi293F cells and purified by Ni-NTA affinity and size-exclusion chromatography, with the tag retained throughout. A 4,576-compound library was screened at 100 µM against CHI3L1 amine-coupled to a CM5 sensor chip on a Biacore 8K instrument, against a reference surface prepared identically without protein. Compounds exceeding 80% fractional occupancy under a 1:1 binding assumption were classified as primary hits and triaged by sensorgram magnitude and shape, association and dissociation behavior, residual binding and reference-surface response. Surviving compounds were re-evaluated by single-cycle kinetics over a concentration series, with affinities reported as apparent KD from 1:1 fits. The primary screen was performed once; LZ6 was measured in three independent experiments and its KD is reported as individual values and as the mean ± SD. Purification, immobilization, injection and regeneration conditions are given in Supplementary Methods.

### Crystallography

CHI3L1 was incubated with LZ6 at a 1:3 molar ratio and crystallized at 289 K by sitting-drop vapor diffusion against 0.1 M HEPES, pH 7.0, 10% (w/v) PEG 6000. Data were collected at 100 K on beamline 17-ID-1 at the National Synchrotron Light Source II, processed with autoPROC and AIMLESS,[24,25] phased by molecular replacement in PHASER using PDB 8R41,[26] and rebuilt and refined in Coot and PHENIX.[27,28] Coordinates and structure factors are deposited under PDB accession 10JZ; statistics are given in Table S1.

### Ternary complex formation

Partner interactions were measured by TR-FRET in reactions containing 20 nM His-CHI3L1, terbium-labelled anti-His antibody, 100 nM biotinylated Gal3BP or galectin-3 and streptavidin-XL665, with LZ6 at 0.03-10 µM and signal normalized to plate-matched vehicle across five independent runs; donor-only, acceptor-only, autofluorescence and unrelated-His-protein controls were run in parallel, with deviations above 15% classified as interference. Complex formation was confirmed orthogonally by BLI (Gal3BP on streptavidin biosensors, 0.25 µM CHI3L1, LZ6 0-10 µM), by native MS in ammonium acetate with the ternary fraction integrated as ternary signal over all assigned complex signal, and by SEC-MALS with in-line light-scattering and refractive-index detection. Stoichiometry was assessed at CHI3L1:partner ratios of 1:1, 1:2 and 2:1 across LZ6 concentrations. For order-of-addition tests LZ6 was added 10 min before or after partner and equilibrated for 5, 30 or 120 min; cooperativity titrations were fit to null, competitive and positive-cooperativity models and compared by Akaike information criterion across six independent protein preparations. Interface formation was tested by HDX-MS, with complexes labelled in D₂O at 4 °C for 10-600 s and digested online with pepsin, and by crosslinking-MS with disuccinimidyl suberate and tryptic digestion, interprotein crosslinks filtered at 1% FDR; four independent preparations were analyzed per condition by each method, and protection is expressed relative to vehicle. Buffers, instrument settings and software are given in Supplementary Methods.

### Cell models and tumor-cell assays

Five patient-derived GBM models (ATCC HCMI PDM-17, PDM-20, PDM-23, PDM-121, PDM-123; designated GBM-PD01-05), GL261 (Cytion) and normal human astrocytes (ScienCell) were cultured in supplier-recommended media at 37 °C in 5% CO₂, authenticated by STR profiling on receipt and every 6 months, and tested monthly for mycoplasma. For spheroid assays, cells were aggregated 72 h in ultra-low-attachment plates, with CD14⁺ monocytes from three donors added where indicated, then treated with LZ6 at 0.01-30 µM for a further 120 h; viability (CellTiter-Glo 3D), LDH release and post-washout sphere formation were measured in separate matched wells across three independent runs. Migration and invasion were assessed in 8-µm-pore inserts, uncoated or Matrigel-coated, with LZ6 at 0-10 µM in both chambers for 18 h and a matched viability assay at each concentration. For CRISPR interference, cells were transduced with either of two promoter-targeting guides for CHI3L1 or LGALS3BP or a non-targeting guide; rescue populations additionally received a constitutively expressed wild-type LGALS3BP coding-sequence construct lacking the targeted promoter region and therefore refractory to interference. Knockdown and rescue were confirmed by qPCR and secreted Gal3BP.

### Myeloid assays

CD14⁺ monocytes from eight independent healthy donors (STEMCELL Technologies) were differentiated for 7 days with M-CSF and treated with LZ6 at 0-5.71 µM for 24 h in myeloid-only medium, GBM-conditioned medium or direct co-culture. Phagocytosis of pHrodo Red-labelled apoptotic GBM targets, surface CD11b, CD206, CD86 and HLA-DR, secreted TNF, IL-10, IL-6, CXCL10 and TGF-β by ELISA, and MRC1, IL10, IFIT1, STAT1 and TNF transcripts in sorted CD45⁺CD11b⁺ cells were measured, with at least 20,000 viable events per flow sample. Each donor was an independent biological replicate and technical wells were averaged within donor before analysis. Antibody panels and gating strategy are given in Supplementary Methods.

### In vitro DMPK and safety

Solubility, logD7.4, plasma stability, microsomal and hepatocyte intrinsic clearance, plasma and tissue unbound fractions, PAMPA-BBB permeability, bidirectional MDCK-MDR1 and MDCK-BCRP transport with and without selective inhibitors, cytochrome P450 inhibition across five isoforms and hERG current inhibition were determined in standard assay formats. Conditions are given in Supplementary Methods.

### Animal studies

All procedures were approved by the Weill Cornell Medicine IACUC (protocol 2024-0008) and are reported in accordance with ARRIVE. Female C57BL/6J mice aged 8-10 weeks received LZ6 orally at 3, 10 or 30 mg/kg in 0.5% methylcellulose/0.1% Tween-80, or 2 mg/kg intravenously, with plasma sampled at eight timepoints to 24 h (three mice per dose and timepoint). For brain distribution, tumor-bearing mice received 10 mg/kg orally and were sampled at 1, 2, 4 or 8 h; normal brain and tumor were dissected separately and LZ6 quantified by LC-MS/MS against matrix-matched calibration curves. Unbound fractions were measured by equilibrium dialysis and Kp,uu calculated from matched total and unbound ratios. For efficacy, mice were implanted with 5 × 10⁴ GL261-CHI3L1 cells in the right striatum (2.0 mm lateral, 1.0 mm anterior to bregma, 3.0 mm below dura) and on day 5 randomized to vehicle plus isotype, LZ6 plus isotype, vehicle plus anti-PD-1 or LZ6 plus anti-PD-1 (45 per group, 180 total). LZ6 was given orally at 10 mg/kg daily and anti-PD-1 (RMP1-14) or rat IgG2a at 200 µg intraperitoneally on days 5, 8, 11, 14 and 17. Survival was monitored to humane endpoint by an assessor blinded. Separate satellite cohorts (10 mice per treatment and phase) were sampled at trough and at peak, 4 h after dosing, for tumor drug concentration, complex indices, total CHI3L1, immune profiling, soluble mediators and transcripts; the combination arm was not profiled. Tolerability was assessed over 14 days of daily dosing.

### Statistical analysis

Significance was assessed at a two-sided α of 0.05. The biological unit was the donor for myeloid experiments, the model for tumor-cell experiments, the protein preparation for biophysical experiments and the animal in vivo; technical replicates were averaged within biological unit before analysis. Comparisons used Welch or paired t tests as appropriate with Benjamini–Hochberg adjustment within endpoint families, the size of each family being stated in the corresponding figure legend. Survival was analyzed by Kaplan-Meier estimation with log-rank comparisons, and treatment effects estimated in a 2 × 2 factorial Cox model containing both main effects and their interaction, with proportional hazards tested per term and globally by scaled Schoenfeld residuals. Analyses used R 4.3.3.

## Ethics

All animal studies were performed in accordance with procedures approved by the Weill Cornell Medicine Institutional Animal Care and Use Committee (Protocol # 2024-0008). Reporting follows the ARRIVE guidelines.

## Authorship

Conceptualization: MTG. Methodology: SU, LZ, MT. Investigation: SU, LZ, MT. Formal analysis: SU, LZ, MTG. Structural analysis: MT, SU. Writing-original draft: SU, MTG. Writing-review and editing: all authors. Supervision: MTG. Funding acquisition: MTG.

## Data availability

All data supporting the conclusions are provided in the manuscript and the Supplementary Information. Coordinates and structure factors for the CHI3L1-LZ6 complex are deposited in the Protein Data Bank under accession 10JZ. The human analyses use publicly available data: TCGA-GBM from the Broad Institute GDAC Firehose (stddata 2016_01_28, https://gdac.broadinstitute.org), originally generated by The Cancer Genome Atlas Research Network, and single-cell RNA sequencing from Gene Expression Omnibus accession GSE182109. No new human data were generated.

