## Supporting Information for "Stabilizing the CHI3L1-Gal3BP complex remodels the glioblastoma immune microenvironment and synergizes with PD-1 blockade"

***Supplementary Methods***

*This section provides extended detail for the procedures summarized in the Materials and Methods of the main manuscript. Headings correspond.*

***Human cohort analyses***

TCGA-GBM expression and clinical data were obtained from the Broad Institute GDAC Firehose standardised data run stddata_2016_01_28: Merge_transcriptome agilentg4502a_07_1 and 07_2 (UNC, log2 lowess normalised gene level), Merge_rnaseqv2 illuminahiseq_rnaseqv2 (UNC, RSEM genes normalised, log2(x+1) transformed), Clinical_Pick_Tier1 Level 4, and Mutation_Packager_Oncotated_Calls. Analysis was restricted to primary tumor samples (barcode sample type 01), retaining the first aliquot where a patient contributed more than one. IDH status was derived from non-silent IDH1 or IDH2 calls and IDH-mutant cases were excluded. Overall survival was days to death for deceased patients and days to last follow-up otherwise. CHI3L1 was standardized within cohort and modelled continuously by Cox proportional hazards; cohort estimates were combined by DerSimonian-Laird random-effects meta-analysis with heterogeneity assessed by Cochran Q and I². Correlations are Pearson coefficients on within-cohort standardized values, combined on the Fisher z scale. The two Agilent-4502A archives share one patient and were combined by meta-analysis rather than pooled; the RNAseqV2 cohort shares 135 of its 153 tumors with the arrays and is therefore reported alongside them rather than as independent validation.

Single-cell data were obtained from GSE182109 as CellRanger count matrices, one region per patient. Cells were retained at 500 or more detected genes, 1,000 or more counts and under 20% mitochondrial reads. Compartments were assigned by the highest mean log-normalized score across canonical lineage marker sets, requiring the leading score to exceed the next by 25%; cells not meeting that margin were left unassigned. Counts were summed within patient and compartment and expressed as counts per million, and compartments with fewer than 50 cells in a patient were excluded. Comparisons used paired t tests on log2 CPM with Benjamini-Hochberg adjustment; the exact Wilcoxon signed-rank test was not used because its two-sided floor at six pairs is P = 0.031. Analyses used R 4.3.3 with survival 3.5.8.

***Protein production, SPR screening and validation***

***Expression and purification.*** The coding sequence for human CHI3L1 (residues 1-383) was inserted into a pCMV-based mammalian expression vector containing a C-terminal HRV 3C protease recognition site and polyhistidine tag. The polyhistidine tag was retained throughout purification and crystallization. Recombinant CHI3L1 was produced by transient transfection of Expi293F suspension cells using the Expi293 Expression System (Thermo Fisher Scientific). Cells were cultured in Expi293 Expression Medium at 37 °C with 8% CO₂ under orbital agitation, following the manufacturer’s recommended conditions. Transfection used the ExpiFectamine 293 Transfection Kit at a plasmid DNA-to-reagent ratio of 1:5, and conditioned medium was collected 5 days after transfection. The supernatant was clarified by centrifugation and loaded onto a manually packed Ni-NTA column pre-equilibrated with 20 mM Tris-HCl, pH 8.0, 150 mM NaCl, 5% (v/v) glycerol and 10 mM imidazole. After washing, CHI3L1 was eluted in the same buffer containing 300 mM imidazole, pooled, and further purified by size-exclusion chromatography in 20 mM Tris-HCl, pH 8.0, 150 mM NaCl and 5% (v/v) glycerol. Purified fractions were pooled and concentrated to approximately 5 mg mL⁻¹.

***Primary SPR screen.*** Screening was performed on a Biacore 8K system (Cytiva, Marlborough, MA, USA). A total of 4,576 compounds from the Discovery Diversity Set (DDS-10; Enamine, Kyiv, Ukraine) were screened for direct binding to recombinant human CHI3L1. His-tagged CHI3L1 (8.0 µg/mL in acetate buffer, pH 5.0) was freshly immobilized on a Series S Sensor Chip CM5 (catalog no. 29104988; Cytiva) by amine coupling using an Amine Coupling Kit (catalog no. BR100050; Cytiva). Immobilization was performed at 10 µL/min for 420 s, followed by blocking of remaining activated groups with ethanolamine. A flow cell subjected to the same activation and ethanolamine-blocking procedure without protein served as the reference surface. Final immobilization levels ranged from 3,000 to 3,600 response units.

Compound stocks were supplied at 10 mM in DMSO. Using a MINI 96 portable electronic pipette (INTEGRA Biosciences, Zizers, Switzerland), 2 µL of each stock was transferred to an intermediate 384-well plate and mixed with 3 µL DMSO to produce a 4 mM intermediate solution; 3 µL of this was transferred to a screening plate containing 117 µL PBS-P+, giving a final compound concentration of 100 µM and 2.5% DMSO. Compounds were injected sequentially over the CHI3L1 and reference surfaces at 30 µL/min for 120 s, followed by a 600-s dissociation phase. After each injection the surface was regenerated with 10 mM sodium acetate and 250 mM NaCl, pH 4.0, at 30 µL/min for 30 s, followed by a 30-s wash with running buffer. PBS-P+ with 2.5% DMSO was the running buffer. The in-house CHI3L1 ligand W1 was injected as a positive control after every 48 screening cycles to monitor surface activity; running buffer alone served as the negative control, and solvent-correction cycles were performed after every 48 injections. Ninety-six compounds were analyzed per channel without reimmobilization.

Reference-subtracted and solvent-corrected sensorgrams were processed in Biacore Insight Evaluation Software (Cytiva). Compound occupancy was calculated under a 1:1 binding assumption at the screening concentration, and compounds exceeding 80% occupancy were classified as primary hits. Their sensorgrams were examined for response magnitude and shape, association behavior, dissociation during the buffer-wash phase, residual binding, reference-surface response and injection-associated irregularities. Compounds showing weak or irregular responses, slow association, substantial residual binding or evidence of nonspecific surface interaction were not advanced. The primary screen was performed once.

***Concentration-dependent validation.*** Candidates were evaluated in a single-cycle SPR format. Increasing concentrations were prepared in PBS-P+ with 2.5% DMSO and injected sequentially over the CHI3L1 and reference surfaces at 30 µL/min, each for 120 s without intervening regeneration, followed by a final 600-s dissociation phase. After each series the surface was regenerated with 10 mM sodium acetate and 250 mM NaCl, pH 4.0, at 30 µL/min for 30 s. Datasets for which solvent correction fell outside the predefined acceptable range were not evaluated further. Binding data were assessed for response magnitude, concentration dependence, saturation, injection artifacts and agreement between experimental and fitted sensorgrams. Sensorgrams were fitted with a 1:1 interaction model where data quality permitted, and affinities are reported as apparent equilibrium dissociation constants.

***Crystallography***

Purified CHI3L1 at 5 mg mL⁻¹ in 20 mM Tris-HCl, pH 8.0, 150 mM NaCl and 5% (v/v) glycerol was incubated with LZ6 at a protein-to-ligand molar ratio of 1:3. Crystallization trials were performed at 289 K by sitting-drop vapor diffusion, combining the complex with reservoir solution containing 0.1 M HEPES, pH 7.0, and 10% (w/v) PEG 6000. Initial crystals were obtained across pH 5.0–7.0 and conditions were optimized for diffraction quality. Data were collected at 100 K on beamline 17-ID-1 at the National Synchrotron Light Source II using a DECTRIS EIGER X 9M detector at an incident wavelength of 0.97934 Å. Images were indexed, integrated and scaled using autoPROC and AIMLESS. The structure was solved by molecular replacement in PHASER using PDB 8R41 as the search model, rebuilt iteratively in Coot and refined in PHENIX version 1.21.1. Data-collection and refinement statistics are given in Table S1; the electrostatic surface of the complex is shown in Supplementary Figure S7.

***Ternary complex formation***

***TR-FRET.*** Interactions were measured in 20-µL reactions containing 20 nM His-CHI3L1 (SinoBiological, 11227-H08H), 5 nM terbium-labelled anti-His antibody, 100 nM biotinylated partner and 20 nM streptavidin-XL665 in 20 mM HEPES, 150 mM NaCl, 0.01% Tween-20 and 0.1% BSA, pH 7.4. Reactions were incubated 60 min at 22 °C in black 384-well plates. LZ6 was tested at 0.03, 0.1, 0.3, 1, 3 and 10 µM; vehicle contained 0.5% DMSO. Readout on a Tecan SPARK used 340-nm excitation, 620- and 665-nm emission, a 100-µs delay and 400-µs integration, with the 665/620 ratio normalized to plate-matched vehicle. Five independent runs were performed with three technical wells per condition. Donor-only, acceptor-only, autofluorescence and unrelated-His-protein controls were tested at 0.1, 1, 10 and 30 µM; deviations greater than 15% were classified as interference.

***BLI.*** Biotinylated Gal3BP at 10 µg/mL was loaded onto streptavidin biosensors for 180 s, equilibrated for 120 s, and exposed to 0.25 µM CHI3L1 with 0, 0.3, 1, 3 or 10 µM LZ6 on an Octet R8 system (Sartorius). Association and dissociation were each recorded for 300 s at 1,000 rpm and 25 °C in 20 mM HEPES, 150 mM NaCl, 0.02% Tween-20 and 0.1% BSA, pH 7.4.

***Native MS.*** Proteins were exchanged into 100 mM ammonium acetate, pH 7.4, and mixed at 2 µM CHI3L1 and 4 µM Gal3BP with 0 or 10 µM LZ6 for 30 min at 22 °C. Samples were infused at 5 µL/min on a SYNAPT G2-Si HDMS Q-TOF (Waters) with 1.5-kV capillary voltage, 150-V sampling cone, 6-mbar source pressure and a 10-30-V collision-energy ramp. Complex fraction was integrated as ternary species signal divided by all assigned complex signal.

***SEC-MALS.*** Samples of 100 µL containing 10 µM CHI3L1, 20 µM Gal3BP and 0 or 30 µM LZ6 were incubated 30 min at 22 °C and separated on a Superdex 200 Increase 10/300 GL column at 0.5 mL/min in 20 mM HEPES, 150 mM NaCl, pH 7.4, using an ÄKTA pure system (Cytiva) coupled to a DAWN HELEOS-II multi-angle light-scattering detector and an Optilab T-rEX refractive-index detector (Wyatt Technology). Data were analyzed in ASTRA.

***Stoichiometry.*** CHI3L1:partner mixing ratios of 1:1, 1:2 and 2:1 were tested with LZ6 at 0, 1, 3 and 10 µM.

***Order of addition and interaction models****.* LZ6 was added either 10 min before or 10 min after partner protein, and mixtures were equilibrated for 5, 30 or 120 min at 22 °C. Cooperativity titrations used 20 nM CHI3L1, 5 nM terbium anti-His antibody, partner concentrations from 0.01 to 3 µM and LZ6 at 0, 1 or 3 µM. Null, competitive and positive-cooperativity models were fit by nonlinear least squares, and Akaike information criterion and root-mean-square error were used to compare fits across six protein preparations.

***HDX-MS.*** CHI3L1 and Gal3BP at 10 µM each were incubated with 0 or 30 µM LZ6 for 30 min at 22 °C. Exchange was initiated by mixing 5 µL sample with 45 µL D₂O buffer (20 mM HEPES, 150 mM NaCl, pD 7.4) at 4 °C for 10, 30, 120 or 600 s, giving final labelling concentrations of 1 µM each protein and 3 µM LZ6 in 90% D₂O; four independent preparations were analyzed. Exchange was quenched with 0.8 M guanidine hydrochloride and 0.8% formic acid, pH 2.5, on ice, and online pepsin digestion at 0 °C preceded LC-MS. Peptides were analyzed by C18 LC-MS on a Waters ACQUITY UPLC M-Class HDX system coupled to a SYNAPT G2-Si Q-TOF; identification and uptake analysis used ProteinLynx Global Server and DynamX. Undeuterated and maximally deuterated controls were included.

***Crosslinking-MS.*** Proteins at 5 µM each with 0 or 15 µM LZ6 were incubated 30 min at 22 °C, then treated with 1 mM disuccinimidyl suberate for 30 min. Four independent preparations were analyzed, with vehicle-matched and no-crosslinker controls. Reactions were quenched with 50 mM Tris, pH 7.5, for 15 min, reduced with 5 mM DTT, alkylated with 15 mM iodoacetamide and digested overnight with trypsin at 1:50 enzyme:protein. Crosslinked peptides were analyzed by LC-MS/MS on the same platform, searched using pLink and filtered at 1% FDR. Crosslink protection was defined as the reduction in normalized interprotein crosslink signal with LZ6 versus vehicle.

***Cell models and tumor-cell assays***

***Cell models.*** Five patient-derived GBM models were obtained from ATCC: HCMI PDM-17, PDM-20, PDM-23, PDM-121 and PDM-123, designated GBM-PD01 through GBM-PD05, cultured in the supplier-recommended medium for each model. Parental GL261 cells were obtained from Cytion (catalog 305225). Normal human astrocytes were obtained from ScienCell (catalog 1800) and maintained in Astrocyte Medium (catalog 1801). Cells were maintained at 37 °C in 5% CO₂. Human cell models were authenticated by STR profiling at receipt and every 6 months; cultures were tested monthly for mycoplasma by PCR. Baseline expression of CHI3L1, LGALS3BP and LGALS3 across models is shown in Supplementary Figure S8.

***Spheroid viability, LDH and sphere formation.*** GBM-PD01-05 cells were seeded at 2,000 cells per well in 100 µL of GBM culture medium in ultra-low-attachment 96-well plates and aggregated for 72 h. Human peripheral-blood CD14⁺ monocytes (STEMCELL Technologies, catalog 70034) from three donors were differentiated for 6 days with 50 ng/mL M-CSF and added to designated wells at 5,000 cells per well. LZ6 was added at 0.01-30 µM with 0.5% DMSO matched across conditions; after 72 h, 50 µL was replaced with 50 µL of 2× compound solution. After 120 h, viability was measured in separate matched wells using CellTiter-Glo 3D (Promega, G9681), with luminescence normalized to vehicle-treated wells. LDH was measured in 50 µL supernatant using the Pierce LDH Cytotoxicity Assay Kit (Thermo Fisher, 88953), including spontaneous-release and maximum-lysis controls, with cytotoxicity calculated per the manufacturer’s instructions. For sphere formation, separate matched spheroids were washed three times, dissociated with Accutase for 8 min at 37 °C, and viable cells counted by trypan-blue exclusion; 500 viable cells were reseeded per well in compound-free medium in ultra-low-attachment plates, and spheres ≥50 µm were counted on day 10, expressed as spheres per 500 cells seeded. Experiments used four technical wells per condition in each of three independent runs.

***Migration and invasion.*** Assays used 24-well plates with 8-µm-pore inserts (Corning). Cells were serum-starved for 4 h, then 5 × 10⁴ cells in 200 µL serum-free medium were added to each upper chamber, with 600 µL medium containing 10% FBS in the lower chamber. For invasion, inserts were coated with 100 µL growth-factor-reduced Matrigel (Corning) diluted 1:8 in cold serum-free medium and polymerized for 45 min at 37 °C. LZ6 at 0, 0.3, 1, 3 or 10 µM or matched vehicle was present in both chambers. After 18 h, non-migrated cells were removed, membranes fixed in 4% paraformaldehyde for 15 min and stained with 0.1% crystal violet for 20 min; bound stain was extracted in 10% acetic acid and absorbance measured at 590 nm. Each condition was tested in triplicate in three independent experiments, with a matched 18-h viability assay at each concentration.

***CRISPR interference.*** Cells were transduced separately with lentiviral vectors encoding either of two promoter-targeting guide RNAs for CHI3L1 or LGALS3BP, or a non-targeting guide, at an MOI of 0.3 in 8 µg/mL polybrene. Stable populations were selected with 1 µg/mL puromycin for 7 days and recovered for 5 days. Knockdown and rescue were confirmed by SYBR Green qPCR using 500 ng RNA, random-hexamer reverse transcription, RPLP0 normalization and the ΔΔCt method, and by measurement of secreted Gal3BP. Each guide was tested in three independent experiments with technical triplicates. Viability was assessed at 0, 1 and 3 µM LZ6 using the spheroid protocol, normalized to the matched non-targeting-guide, vehicle-treated control.

***Myeloid assays***

CD14⁺ monocytes from eight independent healthy donors were isolated using the EasySep Human CD14 Positive Selection Kit II (STEMCELL Technologies) and differentiated for 7 days in RPMI-1640 containing 10% heat-inactivated FBS, 1% GlutaMAX, 1% penicillin-streptomycin and 50 ng/mL M-CSF. On day 7, cells received LZ6 at 0-5.71 µM for 24 h in myeloid-only medium, GBM-conditioned medium or direct co-culture. Conditioned medium was collected after 24 h, clarified at 1,000 × g for 10 min and mixed 1:1 with fresh myeloid medium. Direct co-culture used 1 × 10⁵ myeloid cells and 2 × 10⁴ GBM cells per well.

For phagocytosis, apoptotic GBM targets were labelled with pHrodo Red, exposed to UV-C and allowed to recover for 4 h; targets were added at a 5:1 target-to-myeloid ratio for 2 h at 37 °C. After two cold-PBS washes, cells were stained with viability dye and anti-CD11b and analyzed by flow cytometry. Surface-marker staining used anti-CD11b, CD206, CD86 and HLA-DR for 30 min at 4 °C in darkness, followed by two washes and fixation in 1% paraformaldehyde for 15 min. At least 20,000 viable events were acquired per sample. Supernatants collected at 24 h were assayed in duplicate for TNF, IL-10, IL-6, CXCL10 and TGF-β by ELISA. For transcript analysis, viable CD45⁺CD11b⁺ cells were sorted, 500 ng RNA was reverse-transcribed, and MRC1, IL10, IFIT1, STAT1 and TNF were quantified by qPCR using RPLP0 and HPRT1 as reference genes. Each donor was an independent biological replicate; technical wells were averaged within donor.

***In vitro DMPK and safety***

Kinetic solubility was measured after 2 h at 25 °C in pH 7.4 PBS containing 1% DMSO, followed by centrifugation at 3,000 × g for 20 min and LC-MS/MS measurement of supernatant. Thermodynamic solubility used excess compound equilibrated for 24 h at 25 °C in pH 2.0, 6.5 and 7.4 buffers. LogD7.4 was measured by 2-h shake-flask partitioning between n-octanol and 10 mM phosphate buffer. Plasma stability used 1 µM LZ6 in human, mouse or rat plasma at 37 °C with samples at 0, 15, 30, 60 and 120 min, quenched with three volumes acetonitrile containing internal standard. Microsomal stability used 1 µM LZ6, 0.5 mg/mL pooled microsomes and 1 mM NADPH at 37 °C, sampled at 0, 5, 10, 20, 30 and 45 min. Hepatocyte clearance used 1 µM LZ6 and 0.5 × 10⁶ cryopreserved human hepatocytes per mL at 37 °C, sampled at 0, 15, 30, 60 and 120 min. Plasma, brain and tumor unbound fractions were measured by equilibrium dialysis at 37 °C for 6 h using 10 µM LZ6. PAMPA-BBB was run for 4 h at pH 7.4 with a 10 µM donor concentration. MDCK-MDR1 and MDCK-BCRP transport was measured bidirectionally for 2 h at 10 µM, with or without 1 µM zosuquidar or 0.5 µM Ko143. CYP inhibition used 0.03-30 µM LZ6 and probe substrates for CYP3A4, CYP2D6, CYP2C9, CYP2C19 and CYP1A2. hERG current was measured at 0.3, 1, 3, 10 and 30 µM.

***Animal studies***

***Pharmacokinetics.*** Female C57BL/6J mice aged 8-10 weeks received a single oral dose of LZ6 at 3, 10 or 30 mg/kg by gavage in 0.5% methylcellulose/0.1% Tween-80 at 10 mL/kg. The intravenous reference dose was 2 mg/kg in 10% DMSO, 40% PEG-400 and 50% saline at 5 mL/kg. Plasma was collected at 0.25, 0.5, 1, 2, 4, 8, 12 and 24 h, using three mice per dose and time point. Full profiles, repeat-dose exposure, absolute bioavailability and expanded non-compartmental analysis are shown in Supplementary Figure S11.

***Brain and tumor distribution.*** Tumor-bearing mice received 10 mg/kg LZ6 orally and were sampled at 1, 2, 4 or 8 h, with three mice per time point; blood was collected before euthanasia. Normal brain and tumor were dissected separately, weighed, homogenized 1:3 w/v in PBS and stored at -80 °C. LZ6 was quantified by LC-MS/MS using a C18 column, positive-ion electrospray, multiple-reaction monitoring and a deuterated internal standard. Matrix-matched calibration curves ranged from 1 to 5,000 ng/mL and were fit by quadratic 1/x²-weighted regression; acceptance criteria were ±15% accuracy and precision, or ±20% at the lower limit of quantification. Unbound fractions in plasma, brain and tumor homogenate were measured by equilibrium dialysis at 37 °C for 6 h using 10 µM LZ6. Kp,uu was calculated as the total tissue-to-plasma concentration ratio multiplied by the tissue-to-plasma unbound-fraction ratio at matched time points.

***Orthotopic implantation and treatment****.* Mice were anesthetized with 2% isoflurane and implanted with 5 × 10⁴ GL261-CHI3L1 cells in 1 µL sterile PBS in the right striatum, at 2.0 mm lateral and 1.0 mm anterior to bregma and 3.0 mm below the dura. Injection lasted 3 min and the needle remained in place for 2 min before withdrawal. On day 5, mice were randomized to vehicle plus isotype, LZ6 plus isotype, vehicle plus anti-PD-1, or LZ6 plus anti-PD-1 (45 mice per group; 180 total). LZ6 was administered orally at 10 mg/kg once daily. Anti-mouse PD-1 clone RMP1-14 or rat IgG2a isotype was administered intraperitoneally at 200 µg per mouse on days 5, 8, 11, 14 and 17. Survival was monitored until humane endpoint or study termination; humane endpoints were 20% body-weight loss, inability to reach food or water, severe neurological impairment, or moribund state.

***Pharmacodynamics and tolerability.*** Separate vehicle- and LZ6-treated satellite cohorts were used, with 10 mice per treatment and sampling phase. Following five daily doses, tumors were collected at trough immediately before the next dose, or at peak 4 h after dosing. Tumor drug concentration, CHI3L1-Gal3BP and CHI3L1-galectin-3 complex indices and total CHI3L1 were measured. Tumor immune profiling was performed at peak; soluble mediators and transcripts were measured at both phases. Tumors were processed by 30-min digestion at 37 °C with collagenase IV and DNase I, followed by flow cytometry. Tissue assays included LC-MS/MS, ELISA, BCA protein measurement, flow cytometry and transcript analysis. The combination arm was not included in pharmacodynamic profiling. For tolerability, vehicle- or LZ6-treated mice received 10 mg/kg orally once daily for 14 days, with body weight and clinical score recorded before dosing.

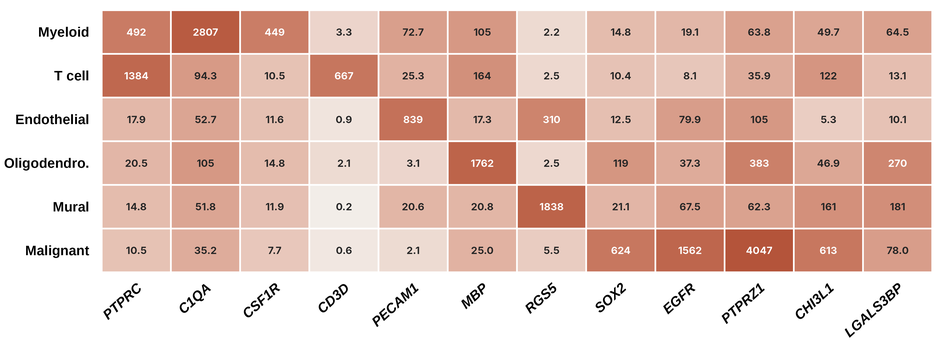

**Figure S1.** Marker validation for single-cell compartment assignment. Pooled expression (counts per million) of canonical lineage markers in each assigned compartment, across the six newly diagnosed glioblastomas analyzed in Figure 1d,e. Compartment labels are not deposited with GSE182109 and were assigned in this work by the markers shown; each lineage marker is enriched in its intended compartment and depleted elsewhere. CHI3L1 and LGALS3BP are included for reference.

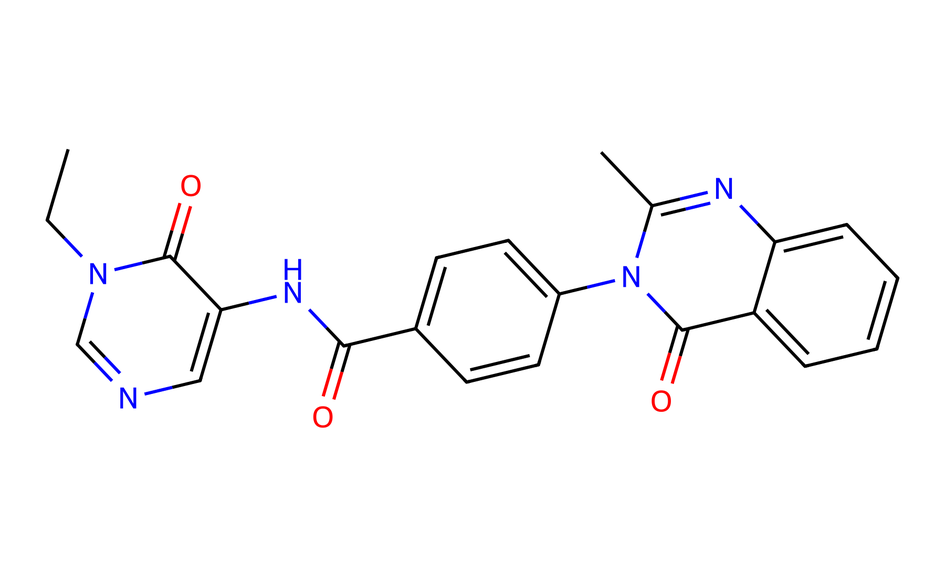

**Figure S2.** Chemical structure of LZ6.

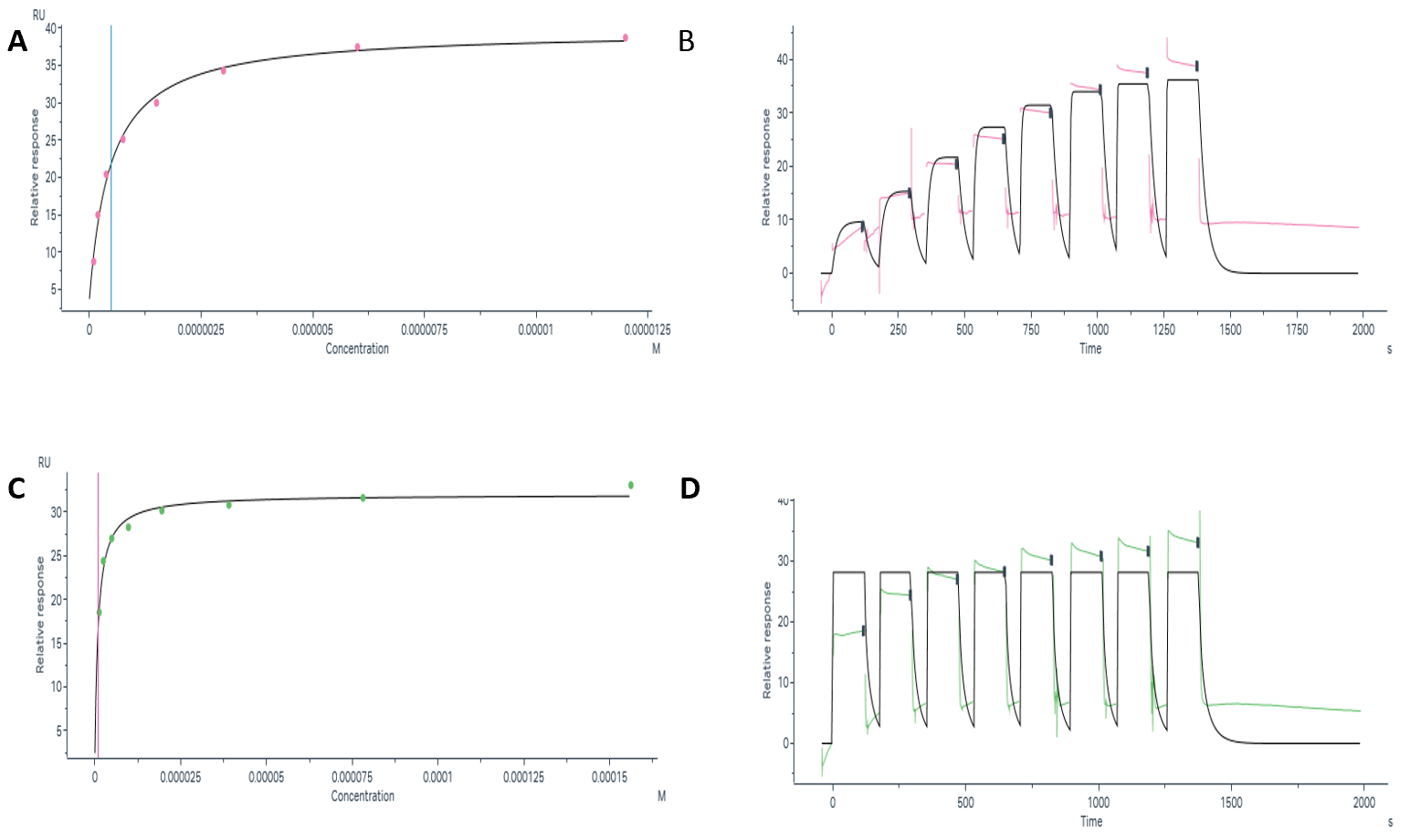

**Figure S3.** Raw binding data. (a-d) Independent surface plasmon resonance measurements of LZ6 binding to immobilized CHI3L1, shown as exported from the instrument software.

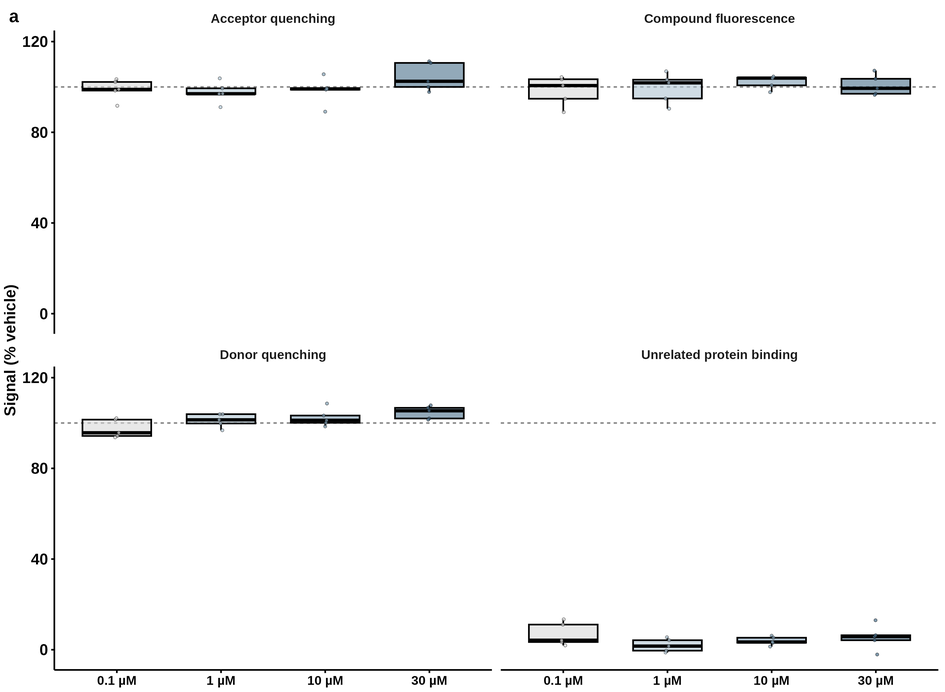

**Figure S4.** Complete assay-interference control set. Donor quenching, acceptor quenching, compound autofluorescence and binding to an unrelated protein at LZ6 concentrations of 0.1, 1, 10 and 30 μM. The dashed line marks vehicle-equivalent signal.

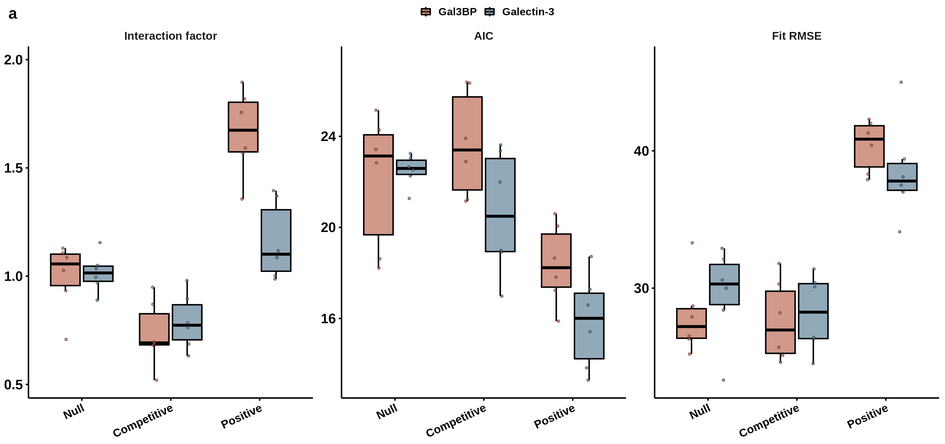

**Figure S5.** Cooperativity model comparison. Fitted interaction factor, Akaike information criterion and fit root-mean-square error for null, competitive and positively cooperative models, for both partner interactions, across six independent protein preparations.

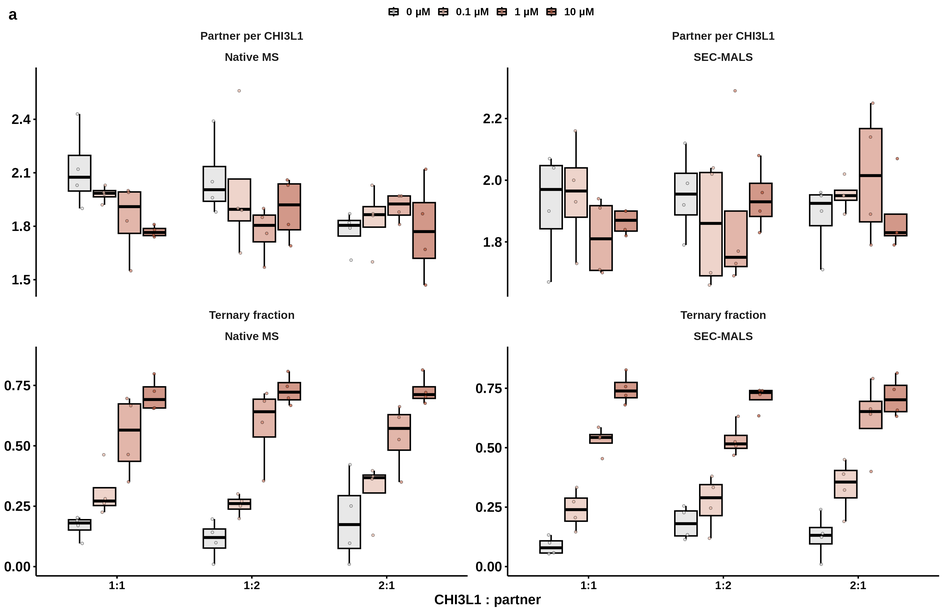

**Figure S6.** Complex stoichiometry and ternary fraction by native mass spectrometry and SEC-MALS at CHI3L1:partner mixing ratios of 1:1, 1:2 and 2:1, across LZ6 concentrations.

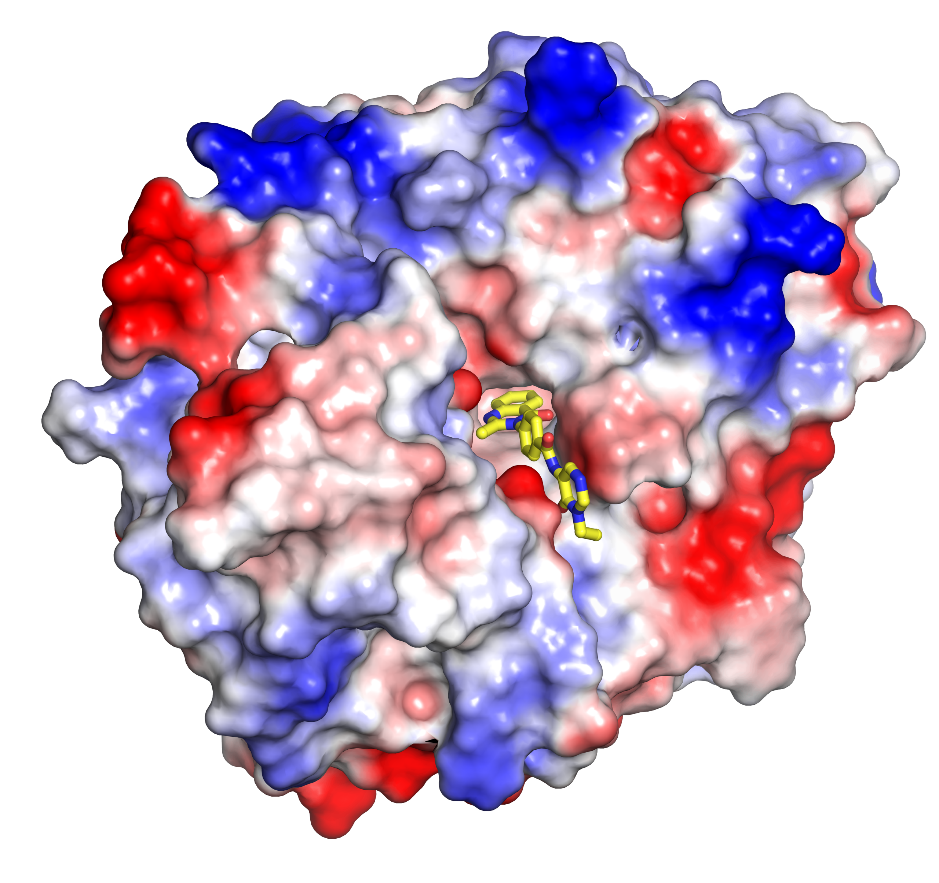

**Figure S7.** Electrostatic surface of the CHI3L1-LZ6 complex. The molecular surface is coloured by electrostatic potential, with negatively charged, neutral and positively charged regions in red, white and blue. LZ6 is shown as yellow sticks. The ligand occupies an elongated, predominantly neutral-to-negatively charged region of the groove, with one end oriented toward the solvent-accessible entrance. PDB 10JZ.

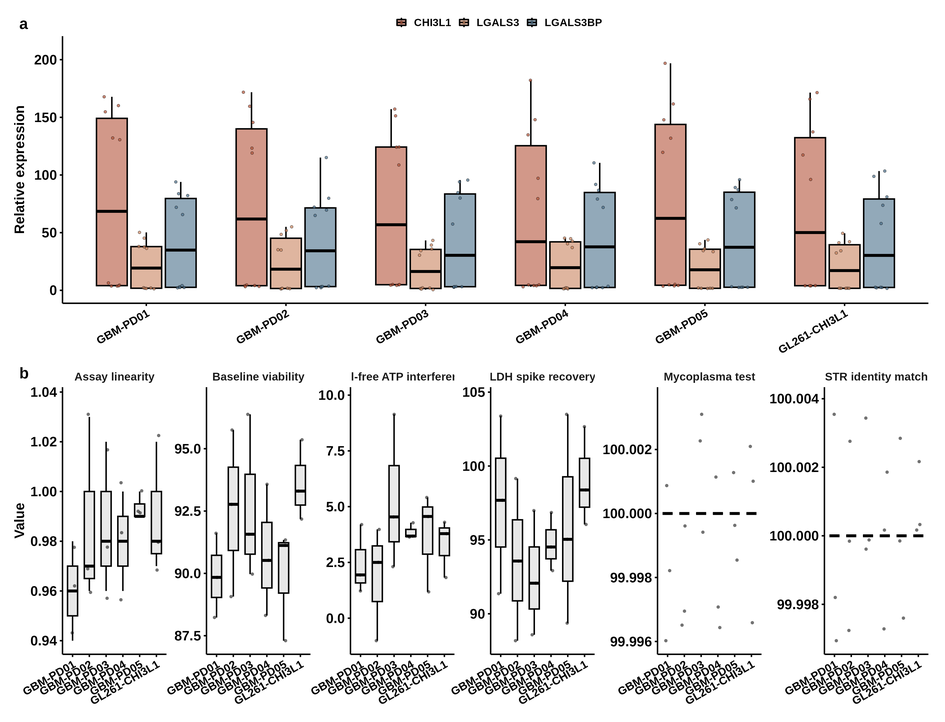

**Figure S8.** Baseline expression of CHI3L1, LGALS3BP and LGALS3 across tumor models.

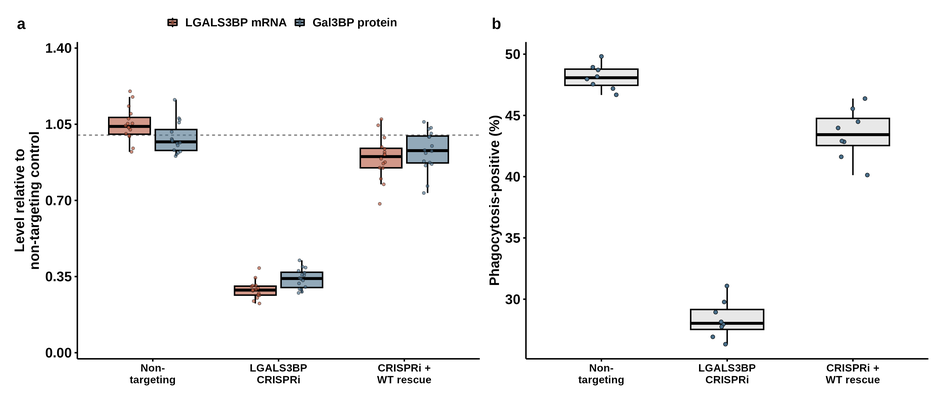

**Figure S9.** CRISPR interference validation. LGALS3BP transcript and secreted Gal3BP protein across guide conditions in eight donors. Interference reduces transcript to 29% and protein to 34% of the non-targeting control, and a CRISPRi-resistant wild-type construct restores both to 90% and 93%; this arm is therefore treated as a successful molecular rescue.

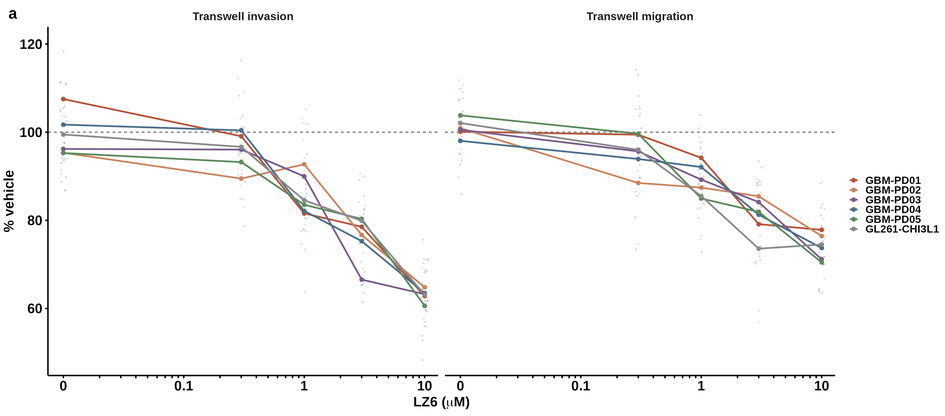

**Figure S10.** Transwell migration and invasion across all tumor models and LZ6 concentrations. Individual replicates are shown with per-model means connected.

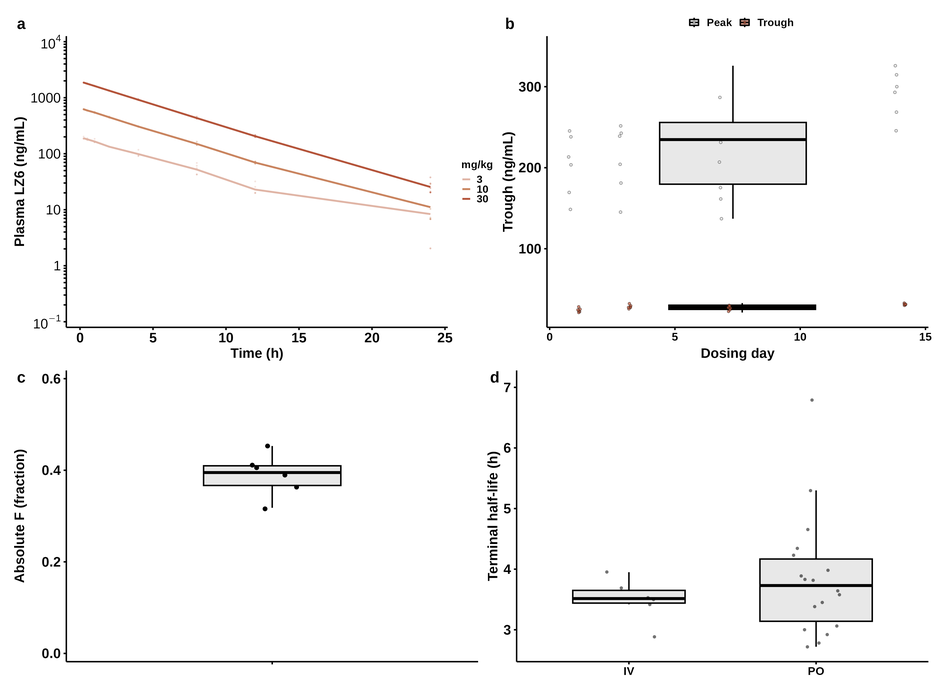

**Figure S11.** (a) Dose-proportionality profiles. (b) Repeat-dose trough exposure. (c) Absolute oral bioavailability. (d) Terminal half-life by route.

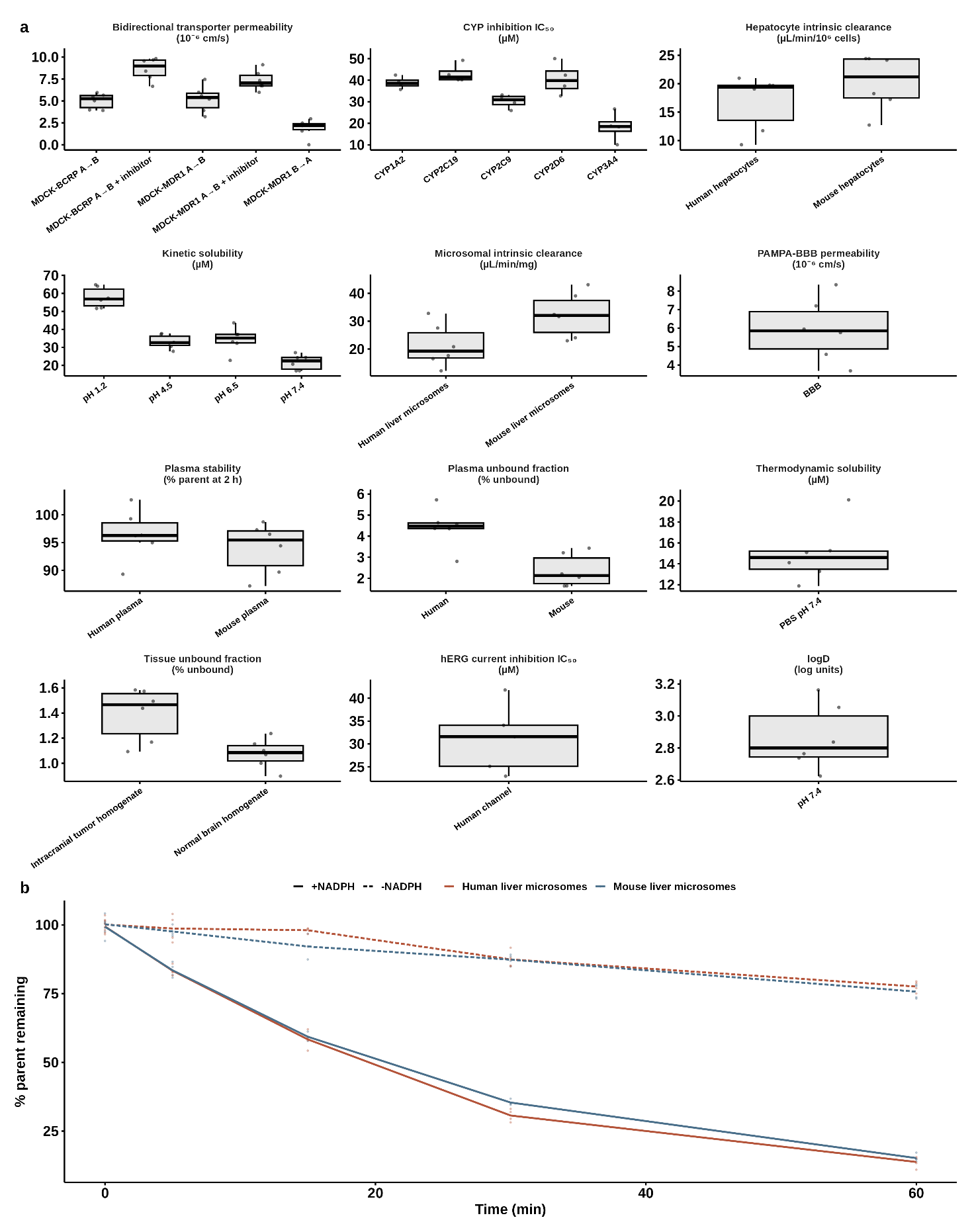

**Figure S12.** (a) Complete in vitro drug metabolism and pharmacokinetics panel: kinetic and thermodynamic solubility, logD, microsomal and hepatocyte intrinsic clearance, plasma stability, plasma and tissue unbound fractions, PAMPA-BBB permeability, bidirectional transporter permeability, cytochrome P450 inhibition and hERG current inhibition. (b) Microsomal time course with and without NADPH.

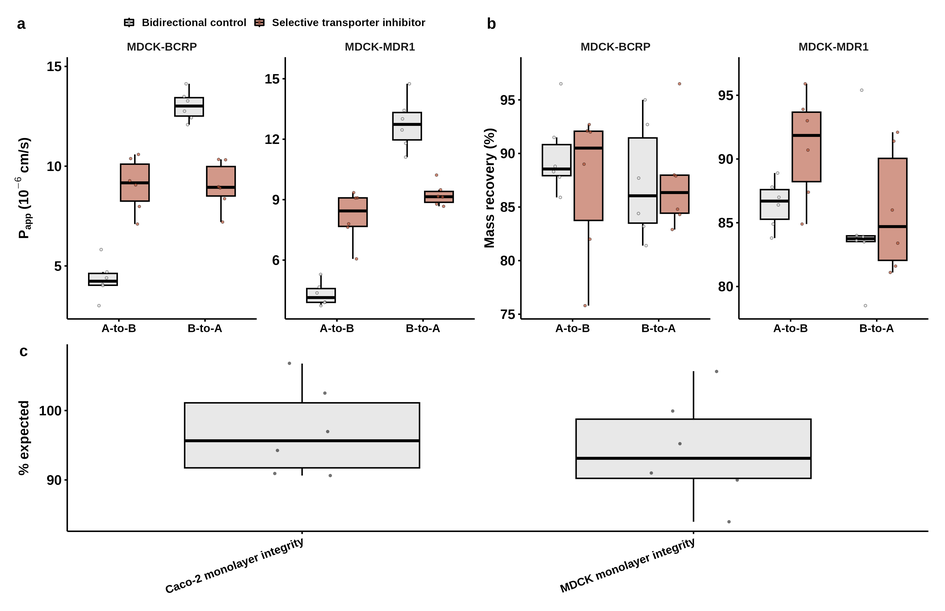

**Figure S13.** (a) Apparent permeability and (b) mass recovery in MDCK-MDR1 and MDCK-BCRP monolayers, bidirectionally, with and without selective transporter inhibitors. (c) Monolayer integrity controls.

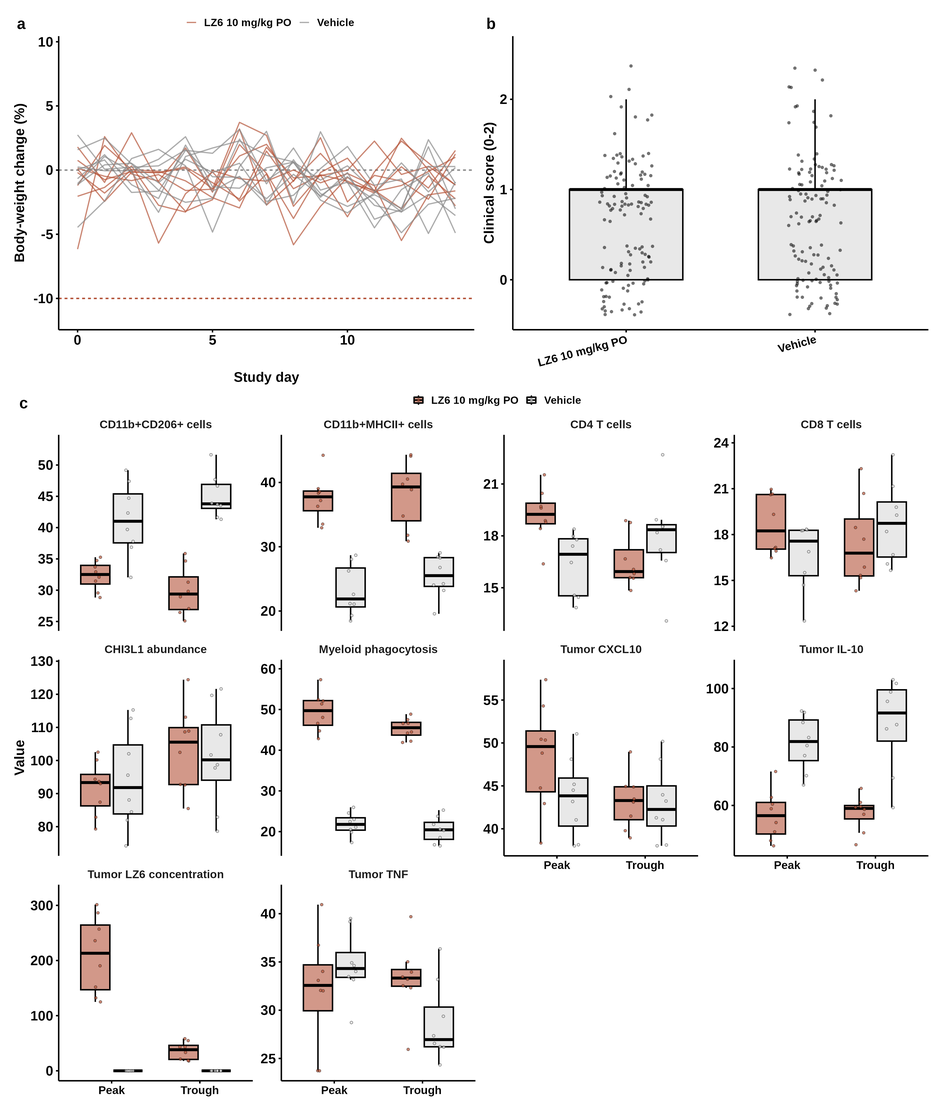

**Figure S14.** (a) Individual body-weight trajectories across fourteen days of dosing; dashed lines mark baseline and a 10% loss threshold. (b) Clinical scores. (c) Satellite pharmacodynamic endpoints at trough and peak sampling, including tumour drug concentration and both partner complex indices.

Supplementary tables

**Table S1.** X-ray data collection and refinement statistics for the CHI3L1-LZ6 complex. Values in parentheses refer to the highest-resolution shell.

*Values in parentheses are for the highest-resolution shell.*

| **Statistic** | **CHI3L1-LZ6** |
| --- | --- |
| **Structure** | |
| PDB accession code | 10JZ |
| Protein molecules in the asymmetric unit | 4 |
| LZ6 molecules in the asymmetric unit | 4 |
| **Data collection** | |
| X-ray source | NSLS-II beamline 17-ID-1 |
| Detector | DECTRIS EIGER X 9M |
| Data-collection temperature (K) | 100 |
| Wavelength (Å) | 0.97934 |
| Space group | P 2₁ 2₁ 2₁ |
| Unit-cell dimensions |  |
| a, b, c (Å) | 109.302, 122.352, 136.250 |
| α, β, γ (°) | 90.00, 90.00, 90.00 |
| Resolution range (Å) | 33.83–2.61 (2.75–2.61) |
| Measured reflections | 245,810 (30,345) |
| Unique reflections | 52,412 (6,779) |
| Completeness (%) | 93.8 (84.3) |
| Multiplicity | 4.7 (4.5) |
| Mean I/σ(I) | 3.7 (1.0) |
| Rmerge | 0.343 (1.911) |
| Rmeas (Rrim) | 0.381 (2.128) |
| Rpim | 0.161 (0.911) |
| CC1/2 | 0.966 (0.400) |
| **Refinement** | |
| Resolution range (Å) | 33.83–2.75 |
| Completeness used in refinement (%) | 95.01 |
| Reflections used in refinement | 45,761 |
| Rwork | 0.1944 |
| Rfree | 0.2390 |
| **Model and validation** | |
| Non-hydrogen atoms | 11,807 |
| Protein | 11,394 |
| LZ6 | 120 |
| Other ligands and glycans | 144 |
| Water | 149 |
| Mean B factor (Å²) |  |
| Protein | 43.72 |
| LZ6 | 44.53 |
| Water | 51.77 |
| R.m.s. deviation in bond lengths (Å) | 0.009 |
| R.m.s. deviation in bond angles (°) | 1.130 |
| Ramachandran favored (%) | 97.70 |
| Ramachandran outliers (%) | 0.21 |
| Rotamer outliers (%) | 1.16 |
| All-atom clashscore | 16.33 |
